# Environmental association identifies candidates for tolerance to low temperature and drought

**DOI:** 10.1101/405399

**Authors:** Li Lei, Ana M. Poets, Chaochih Liu, Skylar R. Wyant, Paul J. Hoffman, Corey K. Carter, Richard M. Trantow, Brian G. Shaw, Xin Li, Gary J. Muehlbauer, Fumiaki Katagiri, Peter L. Morrell

**Affiliations:** Department of Agronomy and Plant Genetics, University of Minnesota, St. Paul, MN 55108; Department of Plant and Microbial Biology, Microbial and Plant Genomics Institute, University of Minnesota, St. Paul, Minnesota 55108

**Keywords:** cold, drought, adaptation, barley, allele frequency differentiation, mixed model association

## Abstract

Barley *(Hordeum vulgare* ssp. *vulgare)* is cultivated from the equator to the Arctic Circle. The wild progenitor species, *Hordeum vulgare ssp. spontaneum*, occupies a relatively narrow latitudinal range (~30 – 40° N) primarily at low elevation (< 1,500 m). Adaptation to the range of cultivation has occurred over ~8,000 years. The genetic basis of this adaptation is amenable to study through environmental association. Using genotyping from 7,864 SNPs in 803 barley landraces, we performed mixed model association analysis relative to bioclimatic variables and analysis of allele frequency differentiation across multiple partitions of the data. Using resequencing data from a subset of these landraces, we tested for linkage disequilibrium (LD) between SNPs queried in genotyping and SNPs in neighboring loci. Six loci previously reported to contribute to adaptive differences in flowering time and abiotic stress in barley and six loci previously identified in other plant species were identified in our analyses. In many cases, patterns of LD are consistent with the causative variant occurring in the immediate vicinity of the queried SNP. The identification of barley orthologs to well characterized genes may provide new understanding of the nature of adaptive variation and could permit a more targeted use of potentially adaptive variants in barley breeding and germplasm improvement.

## Introduction

Cultivated species typically undergo adaptation to very distinct climates as they are disseminated from their region of origin (Gaut et al., 2018). These bouts of adaptation are most extreme for widely cultivated species such as barley and wheat, which are grown from the equator to the Arctic Circle.

Loci that contribute to adaptive phenotypes have typically been identified using top-down approaches (Ross-Ibarra et al., 2007). A phenotype is measured and quantitative trait locus (QTL) mapping or association (also known as linkage disequilibrium or LD) mapping is used to identify genetic variants correlated with the phenotype. Bottom-up approaches that identify genetic evidence of local adaptation or genomic signatures of selection have rarely been used to move from initial analysis to fully characterized genes (Morrell et al., 2012) (for an exception, (see Comadran et al., 2012)).

Genes identified in top-down approaches, by definition, contribute to measurable trait variation, but additional evidence is required to determine if loci identified have played a role in adaptation (Kantar et al., 2017; Ross-Ibarra et al., 2007). For example, the loss of inflorescence shattering in Asian rice was mapped to two loci (Konishi et al., 2006; Li et al., 2006). Only one locus, *qSH1*, shows evidence of selection in both the *japonica* and *indica* subspecies of rice, suggesting that only *qSH1* played a direct role in domestication of both subspecies (Zhang et al., 2009).

Wild barley *(Hordeum vulgare* ssp. *spontaneum)*, the progenitor of cultivated barley *(Hordeum vulgare* ssp. *vulgare)*, occurs primarily within a limited latitudinal range of 30 – 40° N (Harlan and Zohary, 1966). The geographic range of wild barley is bisected by the Zagros Mountains, with peaks of 4,400 m, but wild barley is largely limited to sites at < 1,500 m (Zohary et al., 2012).

Barley was domesticated from its wild progenitor ~10,000 – 12,000 years ago. Domestication occurred at least twice (Morrell and Clegg, 2007) and involved genetic contributions from across the geographic range of wild barley (Poets et al., 2015b). The dissemination of cultivated barley beyond the initial centers of origin began ~2,000 years after domestication (Willcox, 2002). Barley landraces and modern cultivars are the result of pre historic adaptation to growing conditions in Eurasia, North Africa, and much more recently, to Australia and the New World. In Eurasia, the process occurred as humans adopted an agropastoral lifestyle and spread from the Fertile Crescent into a variety of geographic regions. This included cultivation in regions with cooler and more mesic climates in Europe (Pinhasi et al., 2005) as well as drier climates in North Africa and Central Asia (Harris and Gosden, 1996). Barley is frequently produced at high elevations in East Africa, Asia, and Europe and remains among the most important crops in Nepal and Tibet, where it is grown at elevations up to 4,700 m.

The adaptation of cereals such as barley and wheat to northern latitudes or dry climatic conditions involved changes in vernalization requirements (Yan et al., 2006; Dawson et al., 2015), growth habit (Turner et al., 2005; Zakhrabekova et al., 2012; Dawson et al., 2015), and flowering time (Comadran et al., 2012; Dawson et al., 2015). Wild species adapted to Mediterranean climates typically grow over winter and flower in the spring. This is known as a winter growth habit. Under cultivation, winter annuals such as barley and wheat have been adapted to colder climates through spring planting, known as spring growth habit. Spring planting can make cultivation possible at higher latitudes but also increases exposure to frost and freezing conditions (Visioni et al., 2013).

The genetic basis of vernalization and flowering time adaptation in barley has been explored extensively and multiple genes have been cloned (see Hansson et al., 2018). There are also numerous mapping studies and a smaller number of functional studies that have identified regions of the genome associated with cold tolerance in barley (Francia et al., 2004; Hayes et al., 1993; Reinheimer et al., 2004; Skinner et al., 2006; Tondelli et al., 2006; Visioni et al., 2013). Among genes involved in cold tolerance, only the CBF gene cluster has been extensively characterized (Knox et al., 2010), as collecting cold tolerance phenotypes that go beyond “winter survival” is challenging (Visioni et al., 2013). Drought tolerance in barley has also been extensively explored, but the genetic basis of tolerance remains poorly characterized (Honsdorf et al., 2014).

Approaches for the identification of genetic variants contributing to environmental adaptation must discriminate between the effects of selection and neutral evolutionary processes (Rellstab et al., 2015). Demographic effects acting on populations impact the entire genome, whereas selection alters allele frequencies at individual loci (Cavalli-Sforza, 1966). Lewontin and Krakauer (1973) proposed an approach to identify variants subject to differential selection between populations based on allele frequency differences, measured by F-statistics. F-statistic-based comparisons suffer from several weaknesses, including a high expected variance in F_ST_ values (Nei and Maruyama, 1975) and the often arbitrary nature of the partitioning of populations (Lotterhos and Whitlock, 2014). If informative population partitions are defined, F_ST_ measures can identify loci subject to strong differential selection (Beaumont and Balding, 2004). Another approach used to identify the genetic basis of environmental adaptation is mixed-model association analysis. This approach explicitly controls for population structure and treats bioclimatic variables, such as average temperatures and rainfall, as “phenotypes” (Eckert et al., 2010; Yoder et al., 2014; Rellstab et al., 2015).

Here, we present allele frequency (*F*_ST_) outlier and mixed-model association analyses applied to a geographically diverse collection of barley landraces genotyped with the barley 9K Illumina Infinium iSelect Custom Genotyping BeadChip (Comadran et al., 2012) to identify loci potentially involved in cold and drought tolerance. For the *F*ST outlier analysis, we focus on partitions of the sample that distinguish unique growth conditions. These include latitude, elevation, and spring versus winter growth habit. To identify the factors that contribute most to allele frequency differentiation, we also calculated *F*_ST_ for a longitudinal comparison, a contrast reported in previous studies (Morrell and Clegg, 2007; Poets et al., 2015b; Saisho and Purugganan, 2007). We address the following questions: 1) Which of the comparisons explains the largest portion of allele frequency differentiation?, 2) How many previously reported cold temperature and drought tolerance-related loci show evidence of contributing to climatic adaptation?, 3) Do barley orthologs for genes associated with adaptive phenotypes show evidence of contribution to environmental adaptation in our sample?, 4) Given the LD expected in a self-fertilizing species, how frequently are SNPs identified in our analyses in the proximity of potentially causative loci? For this final question, we make use of exome capture resequencing from a sample of 62 landraces drawn from the larger panel. This permits a direct estimate of LD between SNPs identified in our broader panel of accessions and variants in a window surrounding each locus.

We identified a total of six barley genes previously reported to be involved in either cold or drought tolerance, or in flowering time. Furthermore, our analyses identified six additional barley orthologs of genes characterized as contributing to these traits in other plant species. A slight relaxation of the empirical cutoff for outlier *F*_ST_ values identified an additional four characterized barley genes and six orthologs from other plants. Considering both allele frequency outlier and bioclimatic association analyses, we identified 282 barley genes not previously reported to be associated with environmental adaptation. Comparisons of LD between SNPs in genotyping and resequencing suggest that roughly a quarter of the genes we identified on the basis of SNP genotyping are strong candidates for association due to the relatively low gene density in barley.

## Materials and methods

### Plant materials

We use 803 accessions of barley identified as landraces based on passport data from a core collection within the United States Department of Agriculture, National Small Grain Collection (Muñoz-Amatriaín et al., 2014). The 803 individuals were collected from Europe, Asia, and North Africa. These cover the range of barley cultivation in human pre-history (Pinhasi et al., 2005; Poets et al., 2015b; Willcox, 2002). Barley growth habits describe planting times. Spring growth habit is most common, and constitutes 617 (76.8%) accessions of our sample. The balance of the sample includes: 142 (17.7%) winter accessions, 16 (2.0%) facultative accessions that can be planted for spring or winter growth, and 28 accessions (3.5%) of unknown growth habit. Barley can also be divided into the ancestral two-row inflorescence type and the more dense six-row type. Our sample includes 542 (67.5%) accessions of six-row barley, 219 (37.3%) two-row accessions, with the balance of 42 accessions of unknown row type. The reported geographic coordinates for each accession were manually confirmed to identify potentially inaccurate locations, and landraces with highly doubtful locations were filtered out (Table S1). The elevations of collection locations were inferred from the NASA Shuttle Radar Topographic Mission (SRTM) 90 m data (http://www.cgiar-csi.org/) on Oct 7, 2015 using the getData function from R package ‘raster’ (Hijmans et al., 2016).

### Genotyping data

All samples were genotyped using the 7,864 SNPs on the 9K Illumina Infinium iSelect Custom Genotyping BeadChip (Comadran et al., 2012) genotyping platform (henceforth referred to as 9K SNPs). The SNPs are distributed across the seven chromosomes of the diploid barley genome. Because of the relatively large size of the barley genome, the SNP panel includes ~ 1 SNP per 648 kb in the 5.1 Gb genome (Consortium, 2012). For more details on the SNP discovery panel see the description in Comadran et al. (2012; 2015b). Cultivated barley is 99% self-fertilizing (Bothmer, 1992; Wagner and Allard, 1991), and thus the number of unique chromosomes sampled is roughly equal to sample size. The genotyped dataset was filtered for monomorphic SNPs and SNPs with > 20% missingness (Supplemental dataset 1). We culled SNPs in complete LD for comparative analyses, maintaining the SNPs with lower missingness (Supplemental dataset 2).

### Estimating crossover relative to physical distance

We identified the physical position of 9K SNPs relative to the barley reference genome (Mascher et al., 2017) (Supplemental dataset 3; Methods S1). The crossover rate in cM/Mb was estimated using SNP physical positions relative to genetic map positions (Muñoz-Amatriaín et al., 2011). A Python script for this calculation and an R script for Locally Weighted Scatterplot Smoothing (LOESS)(Kono et al., 2018) are included in the project repository https://github.com/MorrellLAB/Env_Assoc.

### Sample differentiation

We estimated the degree of differentiation among individuals by principal components analysis (PCA). PCA was performed using the SmartPCA program from the EIGENSOFT package (Patterson et al., 2006) with SNP data converted from VCF using PLINK 1.90 (Chang et al., 2015).

### Exome resequencing data

We generated exome resequencing from 62 landrace accessions from a randomly chosen subset of landraces in the core collection. This includes 37 six-row spring and 25 two-row spring accessions (Table S2). DNA was extracted from leaf tissue collected from a single plant using a standard 2X CTAB isolation protocol (Saghai-Maroof et al., 1984). The exome resequencing was performed using the NimbleGen exome capture design for barley (Mascher et al., 2017) followed by Illumina 125 bp paired-end resequencing at the University of Kansas Medical Center Genome Sequencing Facility, Kansas City, KS. The data were processed using publicly available software integrated with bash scripts in the ‘sequence_handling’ workflow (PJ et al., 2018). Details are in Methods S1. Variant calling is similar to that reported by Kono et al. (2016), with parameters specified in Methods S1.

### Heterozygosity, SNP diversity, and SNP annotation

Observed heterozygosity was calculated by PLINK 1.90 with the flag ‘--het’. The R package ape (Paradis et al., 2004) was used to calculate average percent pairwise difference (Manhattan distance) between accessions. SNPs in coding and noncoding sequences and in amino acid changing positions within genes were identified using ANNOVAR (Wang et al., 2010) with gene models provided by Mascher et al. (2017) (Supplemental dataset 4).

### Bioclimatic and geographic variables

“WorldClim” bioclimatic data at a resolution of 2.5 minutes were downloaded on 07 Oct 2015 using the getData ‘raster’ R function (Hijmans et al., 2016) in the R statistical language (R Core Team, 2017). The latitude, longitude, elevation, and BIO1 to BIO19 values of the collection locations for each landrace are given in the phenotype data file (Supplemental data 5). Environmental variables can be divided into three categories, geographic factors, temperature, and precipitation. The latitude, longitude, and elevation were classified as geographic factors, BIO1 to BIO11 were classified as temperature, and BIO12 to 19 were classified as precipitation. To identify the relationship among the 22 variables given our sample locations, we performed independent components (ICs) analysis using the icaimax function from ‘ica’ R package (Bell and Sejnowski, 1995). ICs are conceptually similar to principal component summaries of data; however, we found that using the top three ICs appears to capture the cold temperature trend better than using the top three PCs (Table S3). Details of IC interpretation and comparison to Bioclim variables are reported in Methods S1.

### Environmental association mapping

To identify associations between genotypes and environmental variables, we used a mixed linear model (Zhang et al., 2010) implemented in the Genome Association and Prediction Integrated Tool (GAPIT) R package (Lipka et al., 2012). We used the genotyping data to infer the population structure by principal component analysis within GAPIT, and used the first three principal components to control for structure in the mixed linear model. Estimated pairwise distance among samples did not identify individuals with close kinship (95% of comparisons had pairwise distance > 0.43 based on the Manhattan distance between accessions) (Figure S1), and thus kinship was not included in the mixed model. We excluded SNPs with minor allele frequency (MAF) ≤ 0.01 from association analysis. We applied the Benjamini-Hochberg false discovery rate (FDR) correction. We report both adjusted *p*-values and FDR with an FDR threshold ≤0.25.

### *F*_ST_ estimation

To compare allele frequency differentiation in partitions of the data we calculated F-statistics (Wright, 1949) for individual SNPs using the measure of Weir and Cockerham (1984) as implemented in the R package ‘HierFstat’ (de Meeûs and Goudet, 2007). The F_ST_ analysis considered five partitions of the data, which were elevation, high latitude, low latitude, longitude, and growth habit.

The elevation comparison used a threshold of 3,000 m to delineate high elevation accessions. This includes accessions from the European Alps, the Caucasus, Himalayan, Hindu Kush Mountain regions, and the Ethiopian Plateau. Since wild barley typically grows below 1.500 m (Zohary et al., 2012), we also compared the allele frequency at three elevations: below 1.500 m, 1,500m – 3,000m, above 3,000 m.

We compared allele frequencies at two latitudinal ranges: (1) within the wild range of the species (30° N – 40° N) versus landraces at latitudes higher than 40° N (high latitude), and (2) within the wild range of the species (30° N – 40° N) versus landraces at latitudes lower than 30° N (low latitude). High latitude includes the northern extent of the range of wild barley and extends across Eurasia from the Central Iberian Peninsula to the Northern Japanese Archipelago. Low latitude includes the southern extent of the range of wild barley. For landraces, this extends from northwestern Africa to just south of the Japanese Archipelago. We also compared low and high latitude versus the wild range in a single comparison.

For a longitudinal comparison, we divided the sample at 48° E, roughly through the Zagros Mountains, which coincides with the major axis of population structure in wild barley (Fang et al., 2014). The final comparison was spring versus winter growth habit, with assignment based on USDA passport information.

To account for differences in sample size among partitions of the data (Table S4) (Bhatia et al., 2013), we used resampling with equalized sample numbers and 10 iterations without replacement. *F*ST estimates for each SNP were averaged across 10 iterations and outliers were identified at the 99th percentile of the distribution. To calculate the p-value for each F_ST_ value we performed 1,000 permutations. The details can be found in Methods S1.

### Identification of previously reported loci related to cold, drought tolerance, and flowering time

A literature search was used to identify genes previously reported to contribute to plant flowering time and cold or drought tolerance. Google Scholar searches were performed with the terms “cold OR freezing OR drought tolerance” or “flowering time” and “plant” or “barley” (Table S5-7). For each publication with these key words in the title or abstract, we looked for evidence that individual genes reported to contribute to flowering time, cold, or drought tolerance. The protein-coding sequence (CDS) of identified genes were used as the query or subject in BLASTN against the barley high-confidence CDS in May 2016 on the IPK Barley BLAST Server (Mascher et al., 2017). Barley genes and their interval information were extracted if the combined “Score,” “Identity,” “Percentage,” and “Expectation” produced the overall highest ranking and the “Query length” was >100 bp. In the event of identical scores, all highest ranked hits were extracted.

The BEDOPS ‘closest-features’ function (Neph et al., 2012) was used to compare the locations of SNPs and gene intervals. Specifically, if the SNPs were located in the gene interval or 10 kb up- or downstream of the closest genes, we considered those SNPs as identifying the closest gene.

### LD around SNPs

For each 9K SNP identified in environmental association analysis or among *F*_ST_ outliers, we calculated LD with surrounding SNPs called from exome capture resequencing data. We focused on 200 kb windows, 100 kb upstream and downstream of the queried SNP. When the queried SNP was also genotyped by exome capture, this SNP was used for the LD analysis. If the queried SNP was not present in exome capture, we extracted the SNPs called from exome-capture sequencing data surrounding the physical position of the queried SNP. Then we performed the LD analysis using the proximate SNP with a MAF similar to the queried SNP. We filtered out SNPs using a MAF threshold of 1% for all of the SNPs called from the exome-capture resequencing data. For LD analysis, filtering of variants could be anti-conservative, thus for this analysis we removed SNPs with ≥ 50% missing data. We used the R package ‘LDheatmap’ (Shin et al., 2006) to calculate *r*^2^ (Lewontin, 1988).

### Inference of ancestral state

The ancestral state for each SNP from both 9K (Supplemental data 6) and resequencing datasets (Supplemental data 7) was inferred using whole genome resequencing data from *Hordeum murinum* ssp. *glaucum* (Kono et al., 2018) with the programs ANGSD and ANGSD-wrapper (Korneliussen et al., 2014; Durvasula et al., 2016;). We chose *H. murinum* ssp. *glaucum* for ancestral state inference because phylogenetic analyses have placed this diploid species in a clade relatively close to *H. vulgare* (Jakob et al., 2004). Previous comparison of Sanger and exome capture resequencing from the most closely related species, *H. bulbosum*, identified substantial shared polymorphism, resulting in ambiguous ancestral states (Morrell et al., 2013). Methods are detailed in Kono et al. (2018). Both minor and derived allele frequencies were calculated using a Python script.

### Haplotype analysis for individual genes

To assess evidence for functional diversity in the immediate vicinity of SNPs identified in our analysis, we examined haplotype-level diversity in loci that flanked associations. We used exome capture resequencing from the panel described above. Overlapping SNP genotyping was extracted from SNP calls in a variant call format (VCF) file using ‘vcf-intersect’ from vcflib (https://github.com/vcflib/vcflib). Missing genotypes were imputed using PHASE (Stephens et al., 2001; Stephens and Scheet, 2005); PLINK 1.9 (Chang et al., 2015) used to convert the VCF format into PHASE format. Homozygotes were treated as haploid and heterozygotes were treated as diploid samples for haplotype identification.

## Results

### Summary of genotyping and resequencing data

After quality filtering of the genotyping and resequencing data and exclusion of landrace accessions without discrete locality information, our SNP genotyping dataset includes 5,800 SNPs in 784 accessions (Figure S2; Table S1; Supplemental dataset 2). Quality filtering of genotyping data resulted in the removal of 352 SNPs with > 20% missingness. A map of our sampled accessions is shown in Figure S3. The exome resequencing includes 482,714 SNPs in 62 samples (Figure S2; Table S2; Supplemental dataset 8). The site frequency spectrum for SNPs in both panels is shown in Figure S4. Average inbreeding coefficients estimated from SNP genotyping data and exome resequencing data are 0.996 (±0.025) and 0.981 (±0.008) respectively.

### Environmental association and *F*_ST_ outliers

We performed independent component analysis on the 19 bioclimatic variables to identify the subset of climate variables that best summarizes the range of environments occupied by barley landraces (Supplemental dataset 9). We identified 32 SNPs with FDR ≤ 0.25 in environmental association with the first three ICs. Loosening FDR to ≤ 0.3 or ≤ 0.4 identified an additional 45 SNPs, or 77 in total. The ICs constitute a somewhat extreme summary of the bioclimatic variables, as the first three ICs included only eight bioclimatic variables (Table S3). The eight variables are not closely correlated to other bioclimatic variables (Figure S5). Limiting the analysis to ICs potentially excludes some of the bioclimatic signal associated with the remaining variables. Thus we also examined each of the bioclimatic and geographic variables independently. The environmental association with bioclimatic and geographic variables and three ICs identified 155 SNPs in significant associations (with FDR ≤ 0.25) (Figure S6; Table S8).

For both elevation and latitude, we calculated a single *F*_ST_ value with the samples divided into three groups (Table S4; Table S9; Figure S7). We also calculated an *F*_ST_ for low and middle elevations relative to high elevation (Table S4; Table S9; Figure S7). While *F*_ST_ values for pairwise comparisons including many barley genes previously associated with adaptive phenotypes (see below), the single *F*_ST_ values fail to identify these candidate loci. Thus we focused reporting of outlier results on the two level comparisons (Supplemental dataset 10). *F*_ST_ comparisons for elevation, latitude, and growth habit identified 203 outliers (using *F*_ST_ values in the upper 2.5% as the threshold) as the threshold) (Figure 1; Figure S6; Table S10). Considering both the environmental association and *F*_ST_ comparisons, we identified a total of 349 unique SNPs putatively associated with environmental adaptation in our genotyping panel (Figure S6).

**Figure 1:**
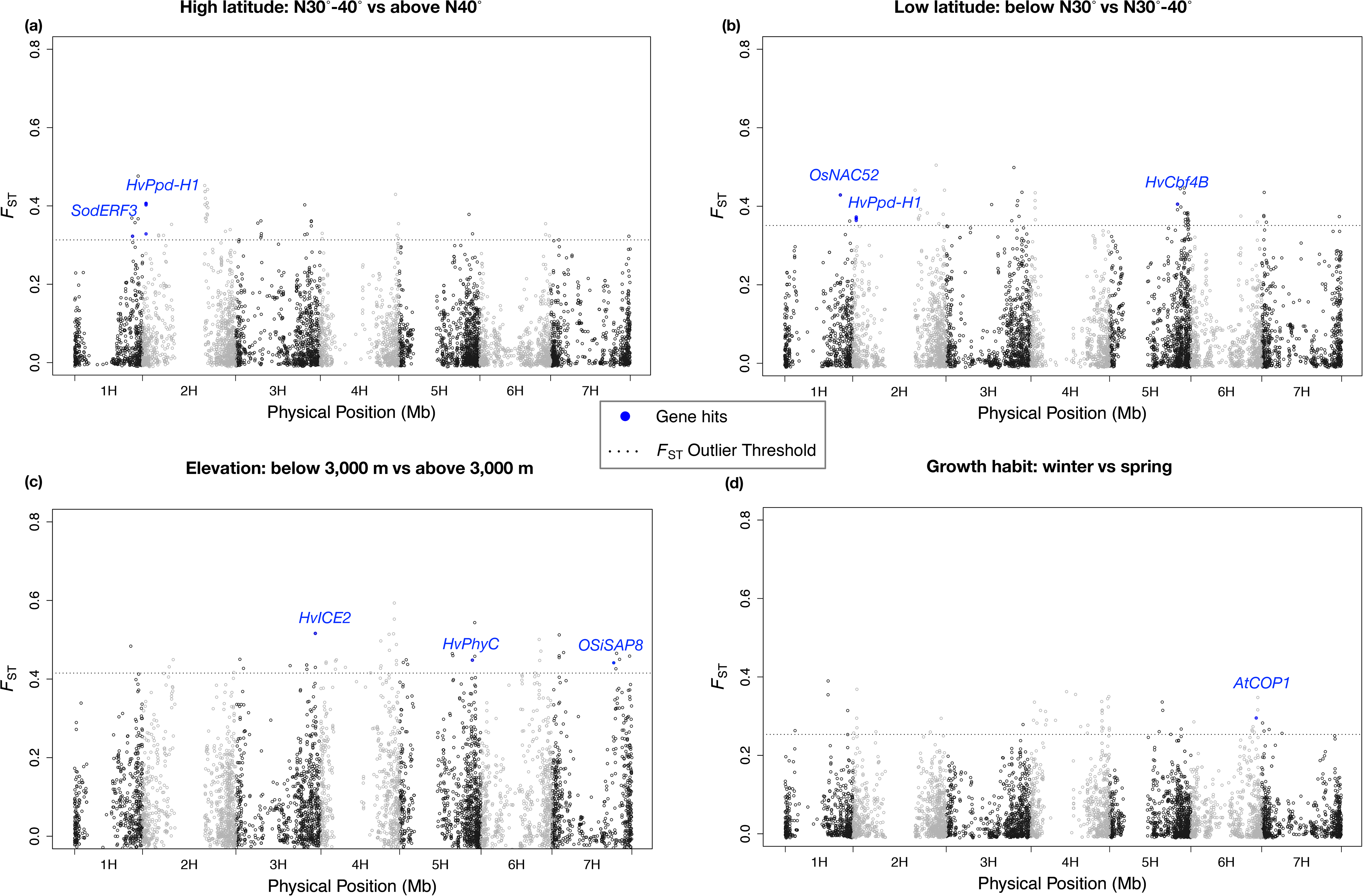
*F*_ST_ for 9K SNPs in samples from comparisons of high latitude (a), low latitude (b), elevation (c), and growth habit (d).

Environmental associations and *F*_ST_ outliers shared nine SNPs in 11 annotated genes. The only characterized gene found in both analyses is *HvPhyC* in barley. For details regarding overlapping results see Table S11.

### Previously reported loci associated with environmental adaptation

Changes in flowering time and drought or cold tolerance are putatively adaptive traits for a cultivated species that has experienced a dramatic expansion in latitudinal range. Our results found four of the 57 genes previously identified in barley as associated with flowering time, two of the 33 genes associated with cold tolerance, and none of the 13 genes associated with drought tolerance (Table 1; Table 3) with the *F*_ST_ threshold set at top 1%. However, we found six genes previously identified in barley as associated with flowering time, four genes associated with cold tolerance, and none of the 13 genes associated with drought tolerance (Table 1), among the 2.5% of *F*_ST_ values.

**Table 1:**
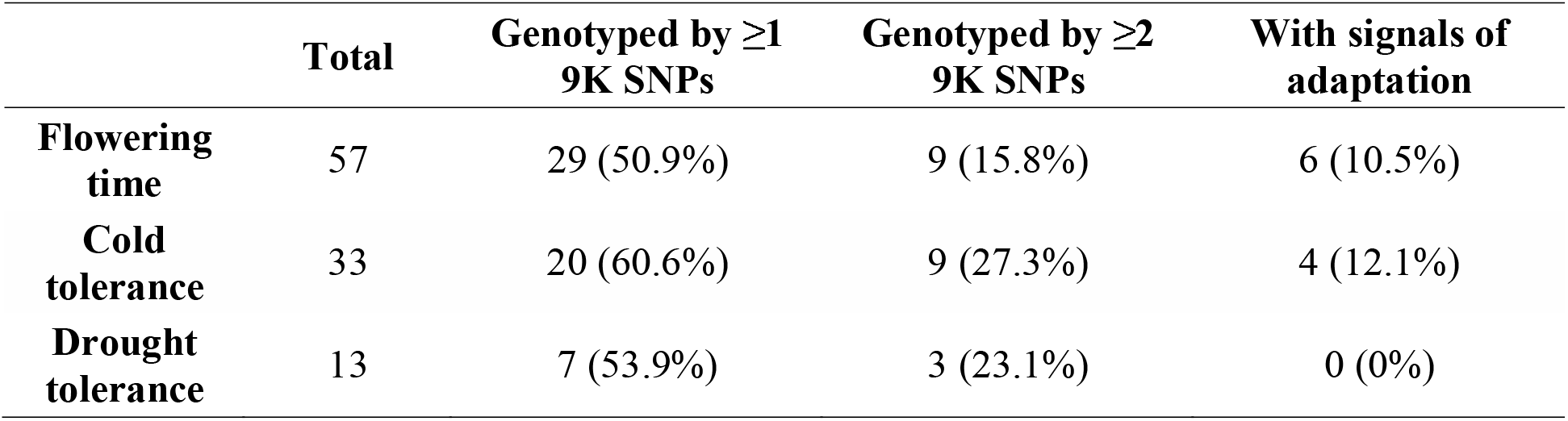
The number of barley genes detected with signals of adaptations and genotyped by 9K SNPs. The number in the parentheses is the fraction of total genes in that functional category that was genotyped or detected with signals of adaptation.

**Table 2:**
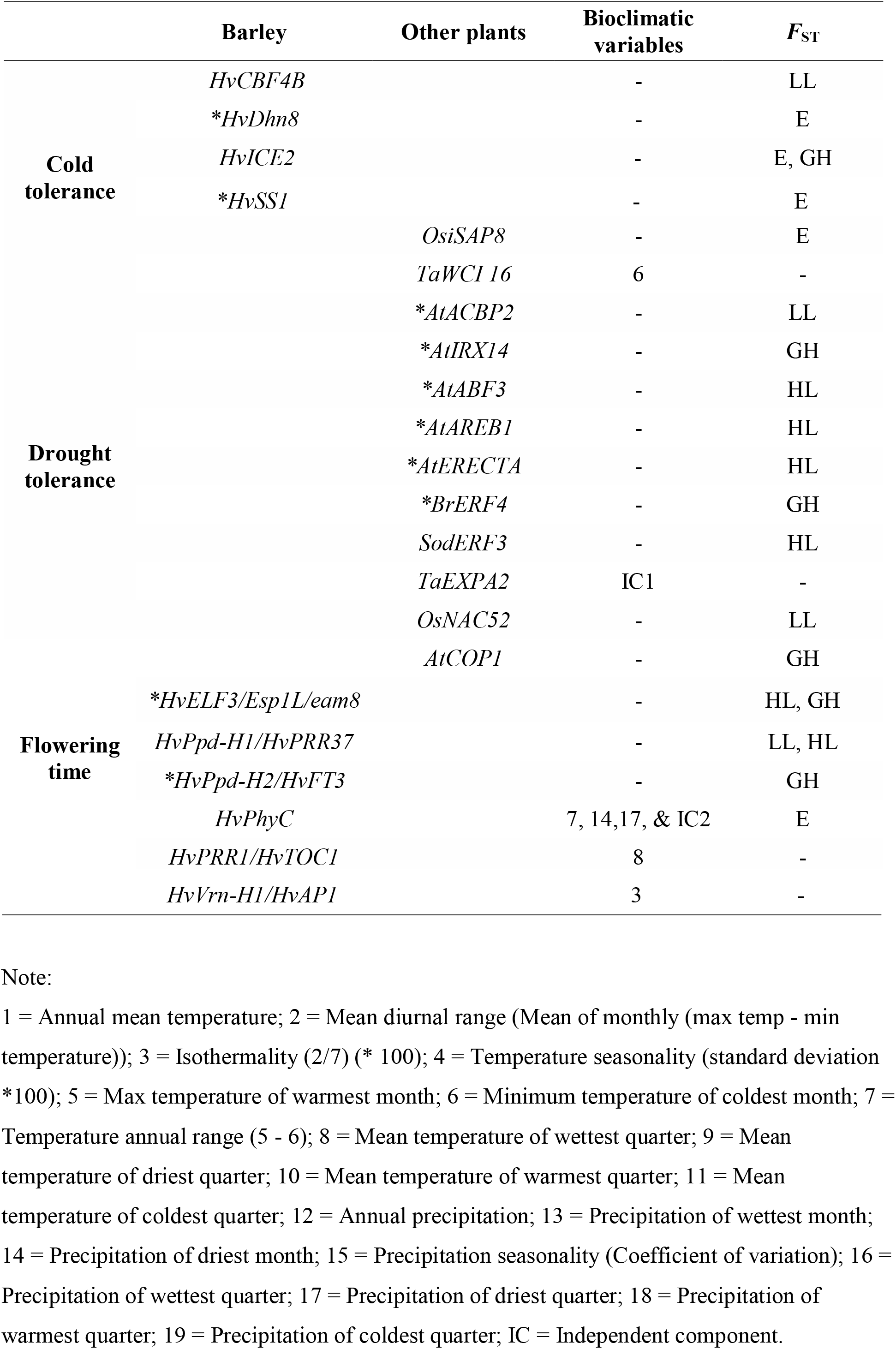
Loci identified in environmental association or *F*ST comparisons that are previously reported to contribute to flowering time, cold, and drought tolerance. Gene names are preceded by two letter prefix with the genus and specific epithet for the species where the gene was identified. This include, *At* – *Arabidopsis thaliana, Ta – Triticum aestivum, Os – Oryza sativa, Br* – *Brassica napus*, and *Sod – Saccharum officinarum.* The *F*_ST_ comparisons involve the following comparisons: E: elevation; LL: Low Latitude; GH: growth habit; HL: high latitude. The * indicates that the gene was identified at the 97.5% threshold but not the 99% threshold.

**Table 3:** The number of SNPs identified by *F*_ST_ outlier approaches and number of previously reported genes they identify. For each comparison, 55 SNPs in total were identified as outliers. Flowering time genes had one across all categories. Drought tolerance had zero SNPs detected in all categories. HL: high latitude; LL: Low Latitude; E: elevation; GH: growth habit.

|  | Flowering time | Cold tolerance |  |
| --- | --- | --- | --- |
|  | SNPs | SNPs | Genes |
| **HL** | 5 | 1 | 1 |
| **LL** | 4 | 2 | 2 |
| **E** | 1 | 1 | 1 |
| **GH** | 1 | 0 | 0 |

The seven loci found associated with flowering time (using *F*_ST_ values in the upper 2.5% of *F*_ST_ values) include four loci identified as *F*_ST_ outliers. *HvPhyC* (Nishida et al., 2013) and *HvPpdH1* (Jones et al., 2008; Turner et al., 2005) occur among the upper 1% of *F*_ST_ values (Table 2; Table 3). *HvELF3 (Esp1L/eam8)* (Boden et al., 2014) and *HvPpd-H2 (HvFT3)* (Casao et al., 2011) are included at the more liberal threshold of *F*_ST_ values in the upper 2.5% (Table 2). Environmental association identified two additional flowering time loci, *HvPRR1 (HvTOC1)* (Ford et al., 2016) and *HvVrn-H1 (HvAP1)* (Cockram et al., 2007), among the 155 outliers at FDR of 0.25 (Table 2).

We also identified four loci previously reported as contributing to cold adaptation in barley, using *F*_ST_ values in the upper 2.5% (Table 3). This includes *HvCbf4B* (Stockinger et al., 2007) and *HvlCE2* (Skinner et al., 2006) as *F*_ST_ outliers for the low latitude, elevation and growth habit comparisons at the top 1% threshold (Table 2). The upper 2.5% threshold for *F*_ST_ includes two additional characterized loci, *HvDhn8* (Choi et al., 1999) and *HvSS1* (Barrero-Sicilia et al., 2011) (Table 2).

A further six loci identified as *F*_ST_ outliers in the top 1% of values or in environmental associations in our barley panel had been previously associated with loci contributing to flowering time or cold or drought tolerance in other plant species (Table 2). This includes one flowering time-related locus characterized in *Arabidopsis thaliana, AtCOP1* (Xu et al., 2016), which was identified as an *F*ST outlier in the top 1% threshold. Two loci (*TaWCI16 and OsiSAP8*) related to cold tolerance were also identified. *TaWCI16* firstly was characterized in wheat and involved in freezing tolerance (Sasaki et al., 2013). The locus was identified in environmental association with “minimum temperature of coldest month (BIO6).” *OsiSAP8* is the rice (*Oryza sativa*) locus, which has been associated with cold, drought, and salt stress response (Kanneganti and Gupta, 2008). *OsiSAP8* was identified by a SNP in the upper 1% of *F*_ST_ values. While no previously identified drought tolerance loci from barley were detected in our analysis, we find evidence of contributions from three loci previously characterized in three other plant species as associated with drought tolerance (Table 2). One of these genes was identified on the basis of environmental association while the other eight involved *F*_ST_ comparisons; only two of the SNPs were included in the upper 1%. In the top 2.5% of values, we found an additional six genes previously characterized in two other plant species as associated with drought tolerance (see Table 2 for details). The identification of multiple characterized loci between upper 2.5% and 1% of *F*_ST_ is indicative of the trade-off between false discovery and false negative rate in empirical scans for adaptive variation (see for example Teshima et al., 2006)

### Relative differentiation among partitions of the sample

Comparison of average *F*_ST_ values provide a means of determining the factors that contribute most to differentiation in barley landraces. Average *F*_ST_ was highest for the longitude comparison with mean genome-wide *F*_ST_ = 0.123 (± 0.13) (Table S9; Supplemental data 10). A primary partitioning of barley populations by longitude, reported as eastern and western populations, has been reported previously (Morrell and Clegg, 2007; Poets et al., 2015b; Saisho and Purugganan, 2007). The second highest average *F*_ST_ was for elevation at 0.089 (± 0.10) (Table S9; Supplemental data 10). The three-level comparison of *F*_ST_ from 0 – 1,500 m, 1,501 – 3,000 m, and >3,000 m resulted in a slightly lower average *F*_ST_ = 0.0826 (± 0.0745) (Table S9; Figure S7; Supplemental data 10). A three-level comparison of latitude with comparisons above and below the range of wild barley (see Materials & Methods for details) was similar to elevation with average *F*_ST_ = 0.087 (±0.082) (Table S9; Figure S7; Supplemental data 10). Pairwise comparisons of the wild range to high latitude, wild to low latitude, and plant growth habit as either spring or winter barley resulted in much lower average *F*_ST_ values (Table S9).

### *F*_ST_ outliers from geographic patterns and growth habit

We focused on comparisons most directly linked to climatic differentiation in *F*_ST_ outliers. We obtained results from the two-level comparisons for high and low latitude, elevation, and growth habit. The upper 1% of *F*_ST_ values from each comparison yielded 55 outlier SNPs for a total of 203 SNPs (Figure 1; Figure S6; Table S10). The comparisons tend to identify unique SNPs. There is overlap of four SNPs in the low and high latitude comparisons and seven SNPs between elevation and growth habit, but other overlaps were not detected (Figure S8). Winter barleys are less frequently grown at higher latitudes and elevations due to harsh winter weather conditions, and indeed winter barleys from these locations are relatively uncommon in the sample, thus constraining the comparisons (Table S1). The elevation comparison identified the largest number of previously characterized loci including *HvPhyC, HvICE2*, and *OSiSAP8* (Figure 1).

SNPs with the most extreme *F*_ST_ values for elevation, growth habit, and latitude comparisons formed very distinctive geographic patterns. Each comparison with the highest *F*_ST_ values occurred with SNPs that fall within genes that are annotated, but uncharacterized. The highest *F*_ST_ from the high latitude comparison occurred at SNP 12_30191 with *F*_ST_ = 0.484 (p-value = 0). The ancestral allele dominates within the wild barley geographic range for this SNP, whereas the derived allele is more prevalent in higher latitude regions (Figure 2a; Supplemental data 6; Supplemental data 10). The highest *F*_ST_ from the low latitude comparison is for the SNP SCRI_RS_153793 with *F*_ST_ = 0.504 (p-value = 0). The ancestral state for the SNP predominates within the geographic range of wild barley and higher latitudes, whereas the derived allele is more prevalent in lower latitudes (Figure S9a; Supplemental data 6; Supplemental data 10). The highest *F*_ST_ between samples from elevation comparison is for SNP 12_20648 with *F*_ST_ = 0.594 (p-value = 0). This SNP’s ancestral allele occurs at high elevations, such as the Himalaya Mountains, while the derived allele tends to occur at lower elevation (Figure 2b; Supplemental data 6; Supplemental data 10). The highest *F*_ST_ between samples from the growth habit comparison was for SNP SCRI_RS_134850 with *F*_ST_ = 0.390 (p-value = 0) (Figure S9b; Supplemental data 6; Supplemental data 10). The SNP SCRI_RS_134850 is significantly negatively correlated with the growth habit with rho = −0.367 (p-value < 2.2 × 10^−16^, Spearman ranking correlation) thus the SNP is predictive of growth habit. The ancestral SNP state is C, and the derived state is T. The CC genotype is observed in 60.5% of winter barley while TT is observed in 88.2% of spring barley (Figure S9b; Supplemental data 6; Supplemental data 10).

**Figure 2:**
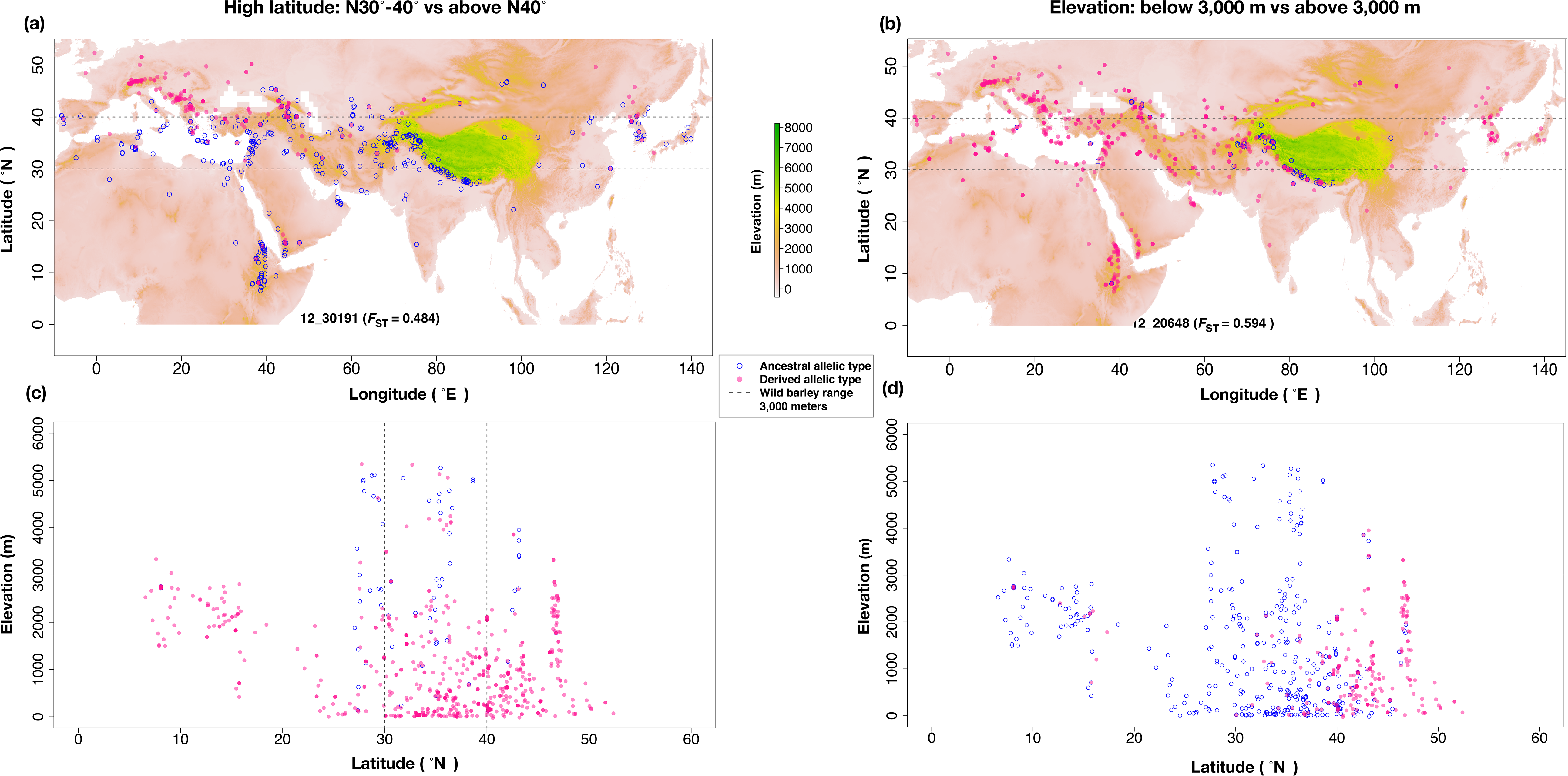
The geographic distribution of the SNPs with high *F*_ST_. (a & c) The geographic distribution of allelic types of 9K SNP 12_30191 with highest *F*_ST_ = 0.4839. The *F*_ST_ was from the high latitude (HL) comparison. (b & d) The geographic distribution of allelic types of 9K SNP 12_11529 with highest *F*_ST_ value of 0.6493. The *F*_ST_ was from elevation (E) comparison. The color bar indicates the elevations in meters. The filled pink circles indicate the derived allele, while the blue open circles indicate the ancestral allele.

### Environmental association to bioclimatic variables

We identified 155 unique SNPs significantly associated with at least one environmental factor with the threshold of FDR < 0.25 (Figure S6; Table S8). All of the p-values and Benjamini-Hochberg FDR-values were reported in Supplemental data 9.

We found 81 SNPs associated with precipitation (variables BIO12 to BIO19) and 51 with temperature (all variables from BIO1 to BIO11) for individual environmental variables (Figure S6). We also identified 47 SNPs associated with geographic variables (latitude, longitude, and elevation), and 32 associated with independent components (top three independent components calculated from BIO1 to BIO19 values after standardization for each BIO variable, called ICs) (Figure S6). Another finding includes 47 cases where individual SNPs were associated with more than one environmental variable (Figure S10). But more generally, as with the *F*_ST_ comparisons, the environmental variables tend to associate with unique sets of SNPs (Figure S10). The largest proportion of unique SNPs were found for precipitation (33.55%), followed by geographic variables (18.71%), temperature variables (18.06%), and then ICs (1.29%) (Figure S10). The aggregated independent components generally did not identify novel variants.

### Minor allele frequency of identified SNPs

The SNPs identified by environmental association and *F*_ST_ have higher MAF than the average MAF across the full SNP data set (0.262 with standard deviation of ± 0.140). SNPs with significant environmental associations have an average MAF = 0.251 (with standard deviation of ± 0.137). The *F*_ST_ outliers have an average MAF = 0.330 (± 0.101). While MAF limits the potential to associate genotype to phenotype for association analysis; the relatively large sample represented here does not suffer from the major limitation of detection. The high MAF contrasts with expectations that adaptive variants for less frequently occupied habitats, such as high elevation sites of cultivation, should be relatively uncommon (average MAF = 0.240 (± 0.091) of outliers from elevation comparison). A relatively low MAF might be expected under models where adaptive variants in a particular environment exhibit antagonistic pleiotropy, and thus confer lower fitness away from habitats in which they are adaptive(Tiffin and Ross-Ibarra, 2014).

### SNP density and LD near focal SNPs

As previously reported, SNP density is highest on chromosome arms and lower in pericentromeric regions (Mascher et al., 2017; Muñoz-Amatriaín et al., 2015). This trend is particularly evident for 9K SNPs (Figure 3b; and Figure S2), and is broadly consistent with lower SNP density in genomic regions with lower observed rates of crossover (Muñoz-Amatriaín et al., 2011). Exome capture density is also lower in pericentromeric regions, such that 51,567 SNPs are detected in 1.560 Gb in pericentromere regions (33 SNPs/Mb) versus 431,147 SNPs in 3.02 Gb (143 SNPs/Mb) on chromosome arms.

**Figure 3:**
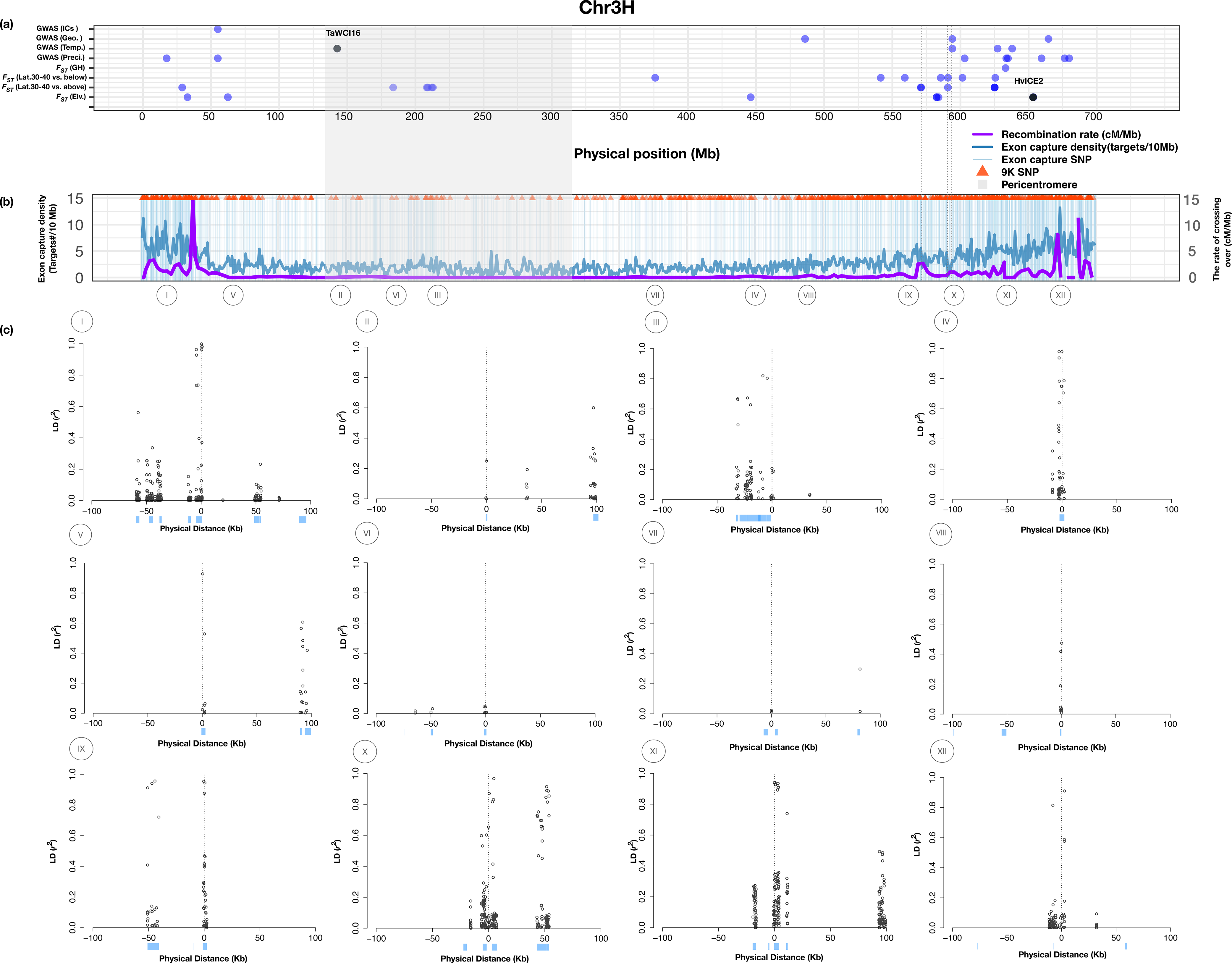
(a) The genomic distribution of outlier SNPs identified according to the *F*_ST_ comparisons of elevation (below 3,000 m vs above 3,000 m), low latitude (below 30°N vs 30-40°N), high latitude (30-40°N vs above 40°N), growth habit (winter vs spring) and association analysis of 21 bioclimatic variables, which are categorized into three classes (precipitation, temperature, and geographic variables) on chromosome 3H; (B) exome capture target density (dark blue line), cross over rate in cM/Mb (purple line), the genomic distribution of SNPs identified in the 62 barley landraces (vertical light blue lines), and 9K SNPs (red triangle) on chromosome 3H; (C) LD plots for SNPs significantly associated with at least one bioclimatic variables (bottom) on chromosome 3H. Each plot shows a 100 Kb window around the SNP. The vertical dotted lines in the upper panel indicate that those outlier SNPs are shared across different traits. For the LD plots, genotyped SNPs are at location 0, and positions upstream and downstream are listed as negative and positive values. The light blue bars are genes in 200 Kb windows surrounding the genotyped SNPs. The SNPs from the I to XII are: 11_20742, 11_10380, SCRI_RS_173916, 12_20108, 11_10601, 12_31008, SCRI_RS_173717, SCRI_RS_6793, SCRI_RS_207408, 12_10210, SCRI_RS_192360, and 12_30960.

We compared LD at queried SNPs to the surrounding region for 358 SNPs identified by environmental association analysis or as *F*_ST_ outliers. We observed 89.3% of these SNPs in exome capture resequencing. The remaining 10.7% of the queried SNPs were replaced by proximal SNPs with similar MAF. The replaced SNPs had an average MAF of 0.035 (±0.005), and were on average 32.9 kb (± 31.5 kb) away from the physical position of the queried SNPs (Figure S11). LD for 123 (34.4%) of these with an *r*^2^ > 0.45 (90th percentile) were limited to SNPs within the same gene (Figure S12a & b; Table 5). Detectable LD with flanking loci is limited in pericentromeric regions because the locus tested is often the only annotated gene within the 200 kb window (Figure 3c). For an additional 212 (59.2%) SNPs, LD extends well beyond the locus where the initial association was identified (Figure S12 c & d). For 23 SNPs (6.4%) there was either no LD with the focal SNPs or no SNPs identified in the 200kb window around the focal SNP (Table 4). These results indicate that the potential to identify individual loci that contribute to adaptive phenotypes is impacted by recombination rate variation and gene density across the genome (Figure 3b, Figure S2).

**Table 4:**
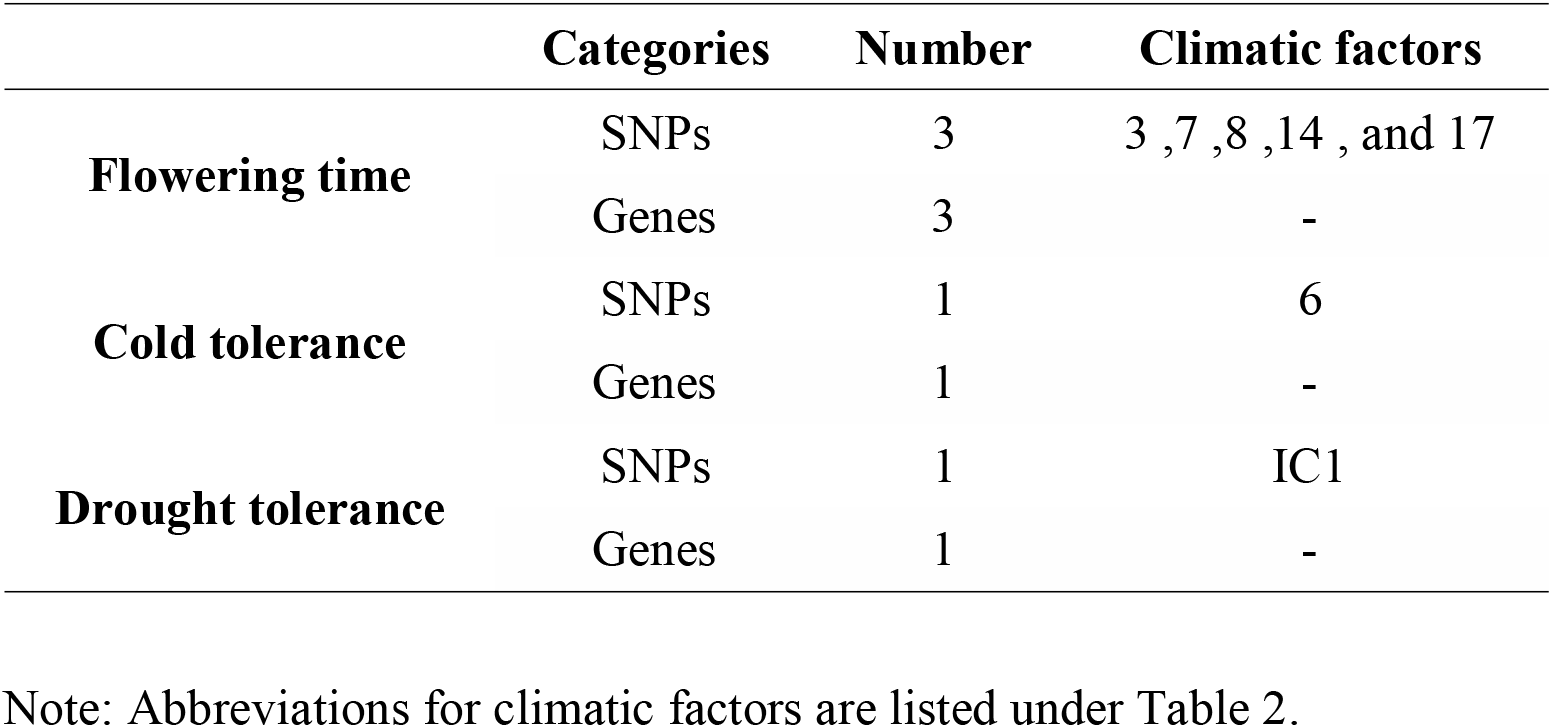
The number of SNPs significantly associated with climatic factors and known genes they hit.

**Table 5:** Linkage disequilibrium for all SNPs associated with environmental variables or identified as *F*_ST_ outliers.

|  | LD within a gene | Extended LD | No LD | Missing* |
| --- | --- | --- | --- | --- |
| $F_{ST}$ | 61 | 132 | 10 | 0 |
| **Association** | 62 | 80 | 7 | 6 |

Note: \* indicates that there are no SNPs in the 200kb window around the target SNP.

### Putative structural variation

An examination of *F*_ST_ outliers prior to LD filtering identified 15 SNPs with *F*_ST_ of ~0.40 for the elevation comparisons. All occur on chromosome 5H at 663.25 cM based on the consensus genetic map(Muñoz-Amatriaín et al., 2011). These SNPs span a physical distance of 133.7 Mb (Table S12). The minor allele frequencies of these SNPs are very similar (0.354 – 0.361) as expected based on *F*_ST_ values, with minor alleles occurring in the same individuals in almost all cases. All 15 SNPs are in nearly complete LD. The region that contains the SNPs is between 131.2 Mb and 265.0 Mb of chromosome 5H, and overlaps with a region identified as a putative chromosomal inversion in wild barley (Fang et al., 2014). The SNPs that Fang et al. (2014) associated with the putative inversion occur between 126.7 Mb and 305.5 Mb. Evidence for an inversion in wild barley was based on elevated *F*_ST_ values, extended LD, and enrichment for environmental associations. Fang et al. (2014) reported a similar pattern on chromosome 2H in wild barley at positions that correspond to 267.3 Mb to 508.7 Mb. We found less evidence of allele frequency differentiation on 2H than in wild barley; observing two SNPs which span ~494 Kb with *F*_ST_ = 0.33 in our sample of landraces (Table S12).

### Haplotype analysis at individual genes

Environmental association results identified a SNP, SCRI_RS_137464, significantly associated with mean “temperature of wettest quarter(BIO8)” (p-value = 8.56 × 10^−4^), which is in the ***HvPRR1/HvTOC1*** gene (Figure 4a). *TOC1* is an important component of the circadian clock in *Arabidopsis.* It conveys crucial function in the integration of light signals to control circadian and morphogenic responses, which is closely related to flowering time (Más et al., 2003). *HvPRR1/HvTOC1* is the ortholog of *TOC1* in ***Arabidopsis thaliana***, and has a high level of sequence similarity and conservation of diurnal and circadian expression patterns when compared to *TOC1* in ***Arabidopsis*** (Campoli et al., 2012). Exome capture resequencing data identified 48 SNPs including SCRI_RS_137464 in ***HvPRR1/HvTOC1***. Five SNPs at the locus annotate as nonsynonymous SNPs and are the most obvious candidates to contribute to functional variation. Four of these are in the last exon of the gene (Figure 4b & c). Five SNPs within ***HvPRR1/HvTOC1*** have relative strong LD with SCRI_RS_137464 (r^2^ >0.45) (Figure 4 a &b). Resequencing identified 20 haplotypes with no obvious geographic pattern (Figure 4c).

**Figure 4:**
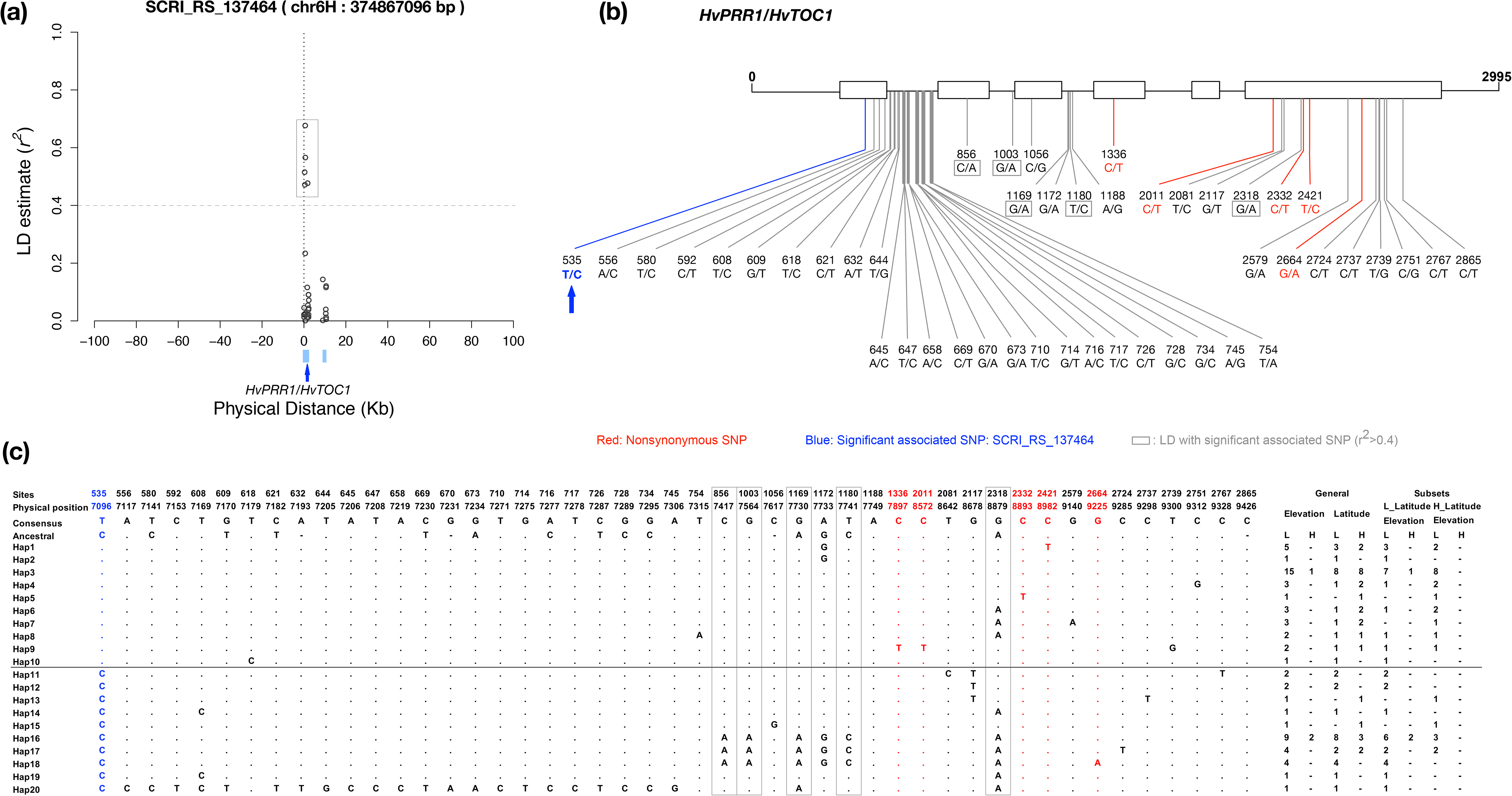
(A) The linkage disequilibrium (LD) analysis of genotyped SNP SCRI_RS_137464 significantly associated with “mean temperature of the wettest quarter (BIO8).” The blue bars indicate genes in the 200 Kb window surrounding SCRI_RS_137464, the red arrow indicates the *HvPRR1/HvTOC1* (flowering time-related gene) that includes SNP SCRI_RS_137464 (B) The gene structure of *HvPRR1/HvTOC1* and the functional annotation of SNPs in this gene. (C) Haplotype structures of *HvPRR1/HvTOC1* based on the SNPs in this gene. L: low; H: high.

Environmental association of both “temperature of coldest month (BIO6)” and “mean temperature of the coldest quarter (BIO11)” identified an association on chromosome 3H with SNP 11_10380 (p-value = 4.95×10^−4^). The SNP is in the barley gene *HORVU3Hr1G030150.1*, which is an ortholog of the wheat gene *WCI16 (Wheat Cold Induced 16)* (Sasaki et al., 2013) (Figure S13). The derived alleles for genotyped SNPs at this locus are much more common in landrace barley than in wild lines. In previous published wild barley genotyping data(Fang et al., 2014) showed the minor allele at 11_10380 occurs in four accessions with geographic provenance information. Those accessions occur at an average of 1460 m – near the upper end of the elevational range for wild barley. Estimated derived allele frequencies differ considerably in wild barley and landraces, at 0.0072 and 0.13 respectively. The 200 Kb window surrounding the SNP contains one gene in addition to *HvWCI16* (Figure S13a). *TaWCI16* encodes a putative transcription factor involved in stomata development. It represents a novel class of late embryogenesis abundant (LEA) proteins in response to cellular dehydration and is involved in freezing tolerance (Sasaki et al., 2013). *TaWCI16* was shown to improve freezing and cold temperature tolerance in wheat when transformed into *Arabidopsis thaliana* (Sasaki et al., 2013). There were six SNPs identified using exome capture sequencing from 61 landraces which includes 11_10380. Three of six SNPs, including 11_10380, are in noncoding sequence (Figure S13b). Of the three SNPs observed in coding regions, one is a nonsynonymous change at nucleotide position 119. This changes from valine to leucine which have similar properties.

There is no evidence of LD between this SNP and others within a 200 Kb window (Figure S13a). Exome capture resequencing identified eight haplotypes, with three of the five being relatively common. Seven haplotypes predominate at lower elevation and lower latitude, with two of those occurring most frequently.(Figure S13c).

Environmental association analysis suggested that the SNP SCRI_RS_235243 significantly (p-value= 3.62×10^−4^) associated with “precipitation of driest months (BIO14)” hit the barley gene ***HORVU1Hr1G008120.1**.* This is an ortholog that produces dehydroascorbate reductase *(**DHAR**;* EC 1.8.5.1) in ***Arabidopsis thaliana*** and bread wheat. It is one of two important enzymes functioning in the regeneration of ascorbate (AsA) which plays a role in protection against oxidative stress (Eltayeb et al., 2006; Osipova et al., 2011). The 200 Kb window surrounding the genotyped SNP SCRI_RS_235243 contains four genes in additional to ***DHAR***, which includes two with exome capture sequence coverage (Figure S14a). Previous results suggest that over expression of ***DHAR*** can protect plants against drought, salt, and polyethylene glycol-induced stress in tobacco and bread wheat (Eltayeb et al., 2006; Osipova et al., 2011). Resequencing identified 53 SNPs in our panel, including SCRI_RS_235243. This encompassed 28 SNPs in noncoding regions, 14 synonymous, and 11 nonsynonymous (Figure S14 b & c). SCRI_RS_235243 is one of nine nonsynonymous SNPs in the first exon of the ***DHAR*** gene (Figure S14 b & c). Six SNPs are in high LD with SCRI_RS_235243 (r^2^ > 0.45), all are noncoding variants within *DHAR* (Figure S14b & c). The derived variant at SCRI_RS_235243 occur within two haplotypes (Figure S14c) that occur in high latitude regions.

A putative causative variant is not immediately apparent for all three of the loci described. However, as the loci *HvPRR1/HvTOC1* and *DHAR* demonstrate, barley landraces are frequently segregating for an abundance of potentially functional variants.

## Discussion

Comparative analysis in a geographically broad collection of Old World barley landraces allowed us to detect environmental associations to bioclimatic variables and identified allele frequency differentiation at six loci with prior evidence of contribution to climatic adaption in barley (Table 1; Table 2). This includes well characterized loci that contribute to flowering time, cold or drought adaptation in barley including ***HvCbf4B*** (Stockinger et al., 2007), *HvICE2* (Skinner et al., 2006), *PhyC* (Nishida et al., 2013), ***HvPpd-H1** (**HvPRR37**)* (Jones et al., 2008; Turner et al., 2005), and *HvVrn-H1 (HvAP1)* (Cockram et al., 2007). All of these loci have been shown to alter phenotypes that are potentially associated with adaptation across the very broad geographic range of cultivation.

### Orthologs that potentially played an adaptive role

We found six loci identified as *F*_ST_ outliers or in environmental associations that had previously been identified as contributing to flowering time, cold or drought stress in other plant species (Table 2). This includes one flowering time-related locus characterized in *Arabidopsis thaliana, AtCOP1* (Xu et al., 2016), which was identified as photomorphogenic repressors and regulates flowering time(Lau and Deng, 2012). Two loci related to cold tolerance were identified. This included wheat locus *TaWCI16* involved in freezing tolerance (Sasaki et al., 2013) and the rice *(Oryza sativa)* locus *OsiSAP8* which has been associated with cold, drought, and salt stress response (Kanneganti and Gupta, 2008). *TaWCI16* was induced during cold acclimation in winter wheat (Sasaki et al., 2013). *OsiSAP8* can be induced by multiple-stresses including heat, cold, salt, desiccation, submergence, wounding, heavy metals, and the stress hormone abscisic acid (Kanneganti and Gupta, 2008). For drought tolerance, we identified nine orthologs characterized in five other plant species (Table 2). For example, over-expression of the wheat expansin gene *TaEXPA2* improved seed production and drought tolerance in transgenic tobacco plants (Chen et al., 2016).

### Why did previously identified genes go undetected in our study?

The genetic basis of flowering time in barley has been explored extensively and multiple genes have been cloned (see Hansson et al., 2018). However, relatively few cold or drought tolerance-related genes have been characterized or cloned (Honsdorf et al., 2014; Visioni et al., 2013). Based on the literature search, we identified 57 flowering time and 33 cold tolerance-related genes in barley (Table S5; Table S6). Our analyses found ~10% of flowering time and ~12% of cold tolerance-related genes (Table 1). We did not identify any of the 13 previously reported drought tolerance-related genes (Table 1).

Why were more previously identified genes not detected? Not every gene was genotyped by the 9K SNPs and many genes genotyped are represented by a single SNP or a small number of SNPs (Table 1). Those SNPs have average MAF of 0.30 (±0.11). As in standard association mapping, the SNPs genotyped need to occur in LD with a causative variant (Balding, 2006); even SNPs in close physical proximity can occur on alternate haplotypes and have limited LD (Nordborg and Tavaré, 2002). For flowering time, 29 of 57 genes were genotyped by at least one SNP. Nine genes were genotyped by two or more SNPs (Table 1). All but one of these genes, *HvVrn-H3/HvFT1*, was found in our analysis (Table S13). All nine of these genes have been identified in previous geographic comparisons in barley (Muñoz-Amatriaín et al., 2014; Russell et al., 2016). Among genes identified by multiple SNPs is *HvPpd-H1/HvPRR37*, a key regulator of flowering (Jones et al., 2008; Turner et al., 2005). This gene was genotyped by eight SNPs, and with five SNPs identified as outliers in our comparisons. Previous studies have identified the *HvCEN* and *HvVrn-H2/HvZCCT-Ha/b/c* genes associated with flowering time as allele frequency outliers (Muñoz-Amatriaín et al., 2014; Russell et al., 2016). *HvCEN* was not genotyped by any SNP in our panel. *HvVrn-H2* was genotyped by a single SNP with an *F*_ST_ = 0.143 in the elevation comparison, at the 75th percentile in this comparison and thus below the 99th percentile threshold. In summary, genes identified through top-down approaches are generally identified in our comparison if they are represented by a sufficient number of SNPs (Table S13). Genes can contribute to trait variation without having played a role in previous rounds of adaptation (Kantar et al., 2017; Ross-Ibarra et al., 2007). However, given current SNP densities, it is premature to conclude that any of the absent loci did not contribute to adaptation in barley.

### Comparison to previous studies

Three of the loci we identified as contributing to adaptive differentiation in Old World landraces were previously reported as *F*_ST_ outliers. They contribute to geographic differentiation in barley breeding populations in in North America (Poets et al., 2015a). This included *HvCbf4B, HvPpd-H1*, and *HvVrn-H1. HvCbf4.* They were also found as an *F*_ST_ outlier and in association with temperature adaptation in comparisons of wild barley populations in the Old World (Fang et al., 2014).

We focused on SNP comparisons, but also found evidence that a large chromosomal inversion has contributed to elevational adaptation in barley. On chromosome 5H, 15 SNPs have *F*_ST_ of ~0.40 in the elevation comparison. All occur at the consensus genetic map position of 663.25 cM. Fang et al. (2014) characterized the region as a putative chromosomal inversion that differs in frequency between the eastern and western portions of the geographic range of wild barley. Recent studies have identified putative chromosomal inversions that contribute to elevation and temperature gradients in teosinte and maize (Fang et al., 2012; Hufford et al., 2012; Pyhäjärvi et al., 2013), rainfall regime and annual versus perennial growth habit in *Mimulus guttattus* (Sweigart and Willis, 2003; Lowry and Willis, 2010), and temperature and precipitation differences in wild barley (Fang et al., 2014). In a close parallel to our results, an inversion on maize chromosome 3 appears to contribute to adaptation of teosinte, which occurs at relatively low elevation to highland cultivation (Wang et al., 2017).

### Advantages of study design and future prospects

SNP density is a limitation in our study. With ~40,000 annotated genes in the barley genome (Mascher et al., 2017), roughly one in six genes was directly genotyped. For roughly one quarter of SNPs, there is limited LD with nearby loci (Figure 3; Figure S12). In regions of the genome with high crossover rates and higher gene density, LD can be limited beyond the locus containing the genotyped SNP (Figure 3c and Figure S2). In regions with limited crossover, gene density is also low (Figure 3b and Figure S2). LD would typically have to extend hundreds of kilobases between genotyped SNPs (Figure S12) and a causative variant at another locus to create an association. High MAF of genotyped variants may also contribute to limited LD. Common variants are typically older and have experienced more recombination, thus they can be closer to linkage equilibrium (Nordborg and Tavaré, 2002).

Our study benefits from largesample size. Russell et al. (2016) performed environmental association with 1,688,807 SNPs from exome capture resequencing in 137 cultivated samples. While the analysis identified 10 loci associated with flowering time, many other previously reported genes went undetected. This prompted the authors to suggest a lack of power owing to small sample size (Russell et al., 2016). Despite limited SNP density and the sampling of relatively common variants, our comparative analysis identified a number of previously identified barley loci and many plausible candidate loci from other plant species. Better coverage of barley gene space through exome capture or whole genome resequencing in a relatively deep panel of accessions would likely uncover a much more comprehensive set of variants contributing to environmental adaptation. This could contribute to targeted use of variants for adaptation to environmental and climatic conditions for barley breeding and germplasm improvement, with the potential to improve the understanding of loci that contribute to climatic adaptation in wheat and other cereals.

## Acknowledgements

We would like to thank A Proulx for annotation of physical positions of 9K SNPs. E Vonderharr assisted with National Center for Biotechnology Information’s Sequence Read Archive (NCBI SRA) submissions. T Kono gave helpful comments to improve the manuscript. This study was supported by the U.S. NSF Plant Genome Program (IOS-1339393) to PLM and the USDA Triticeae Coordinated Agricultural Project 2011-68002-30029 to GJM and PLM, and NSF (MCB-1518058) to FK. This research was carried out with hardware and software support provided by the Minnesota Supercomputing Institute (MSI) at the University of Minnesota.

## Data Accessibility

All sequences were submitted to the NCBI SRA associated with BioProject numbers PRJNA473780 and PRJNA488050. Supporting data (Supplemental data 1-10), including a VCF file of exome capture SNPs and the full results for environmental association and *F*_ST_ are available at https://drive.google.com/drive/folders/1fEwSX2k-7Bbc-wZSWhmM-hCAqmhCcJwQ. Scripts for analysis and associated files are available on a project Github site at https://github.com/MorrellLAB/Env_Assoc.

## Author contributions

PLM, AMP, FK, and LL designed the project. GJM generated the exome-capture resequencing data. RMT and BGS assisted with genotyping data quality control. LL, FK, CL, SRW, CC, XL, and AMP analyzed the data. LL, CL, BGS, and PLM wrote the manuscript.

## Supplemental figures

Methods S1

Figure S1: The distribution of the pairwise genetic distance (Manhattan distance) from 784 barley landraces.

Figure S2: Exome capture target density (dark blue line), crossover rate in cM/Mb (purple line), the genomic distribution of SNPs identified in 62 barley landraces (vertical light blue lines), and 9K iSelect SNPs (red triangle) for seven chromosomes. Crossover rates were calculated using 9K SNPs. SNP genetic positions are based on the genetic map of Muñoz-Amatriaín et al. (2011).

Figure S3: Relationship of barley landrace accessions based on principal components. PC1 and PC2 are depicted relative to a map of the landrace distribution in Africa and Eurasia.

Figure S4: The derived site frequency spectrum (SNPs with inferred ancestral state) for: (a)

2,806 SNPs from the 9K iSelect genotyped in the 784 landraces and (b) 340,260 SNPs with exome-capture resequencing data in 62 landraces. Ancestral state was based on majority state from *H. murinum spp. glaucum* resequencing mapped to the Morex assembly. For all SNPs: (c) Minor allele frequency, for 6,152 SNPs from the 9K iSelect and (d) for 482,714 SNPs from exome-capture resequencing.

Figure S5: The heat map of pairwise correlation coefficient (Pearson correlation) of 22 environmental variables.

Figure S6: The genomic distribution of outlier SNPs identified according to the *F*_ST_ comparisons of elevation (below 3,000 m versus above 3,000 m), low latitude (below 30°N versus 30-40°N), high latitude (30-40°N vs above 40°N), growth habit (winter versus spring) and association analysis of 21 bioclimatic variables, which are categorized into three classes (precipitation, temperature, and geographic variables) and ICs.

Figure S7: The distribution of ranked *F*_ST_ from two- and three-level comparisons of elevation and latitude.

Figure S8: Venn diagram for the *F*_ST_ outliers from the comparisons of elevation, high and low latitude, and growth habit.

Figure S9: The geographic distribution of the SNPs with high *F*_ST_. (a) The geographic distribution of allelic types of 9K SNP SCRI_RS_153793 with highest *F*_ST_ = 0.505. The *F*_ST_ was from the low latitude (LL) comparison. (b) The geographic distribution of allelic types of 9K SCRI_RS_134850 with highest *F*_ST_ = 0.390. The *F*_ST_ was from growth habit (GH) comparison. The color bar indicates the elevations in meters. The filled pink circles indicate the derived allele, while the blue open circles indicate the ancestral allele.

Figure S10: Venn diagram for the candidate SNPs significant associated with three categories of environmental variables: precipitation, temperature, and geographic factors.

Figure S11: The difference between the replaced SNPs and queried SNPs not in the exome-capture data. (a) The minor allele frequency (MAF); (b) The physical distance. The replaced SNPs refer to those SNPs called based on the exome capture resequnecing data and with the similar MAF as the queried SNPs. But the queried SNPs are not in the exome capture resequnecing data.

Figure S12: LD decay plot for 200 Kb window around the significant SNPs associated with environmental variables. The blue bars underneath the x-axis are the annotated genes in the 200 Kb windows. The vertical dashed lines are candidate SNPlocations. The negative signs on the x-axis refer to positions downstream of the candidate SNP.

Figure S13: (a) The linkage disequilibrium (LD) analysis of candidate SNP SCRI_RS_137464 significant associated with “min temperature of coldest month (BIO6)” and “mean temperature of the coldest quarter (BIO6 and 11)”. The blue bars indicate genes in the 200 Kb window surrounding 11_10380, the red arrow indicates the WCI 16 (cold tolerance-related gene) hit by 11_10380, (b) The gene structure of *WCI 16* and the functional annotation of SNPs in this gene. (c) Haplotype structures of *WCI 16*based on the SNPs in this gene. L: low; H: high.

Figure S14: (a) The linkage disequilibrium (LD) analysis of candidate SNP SCRI_RS_235243 significantly associated with ‘‘precipitation of driest months” (BIO14). The blue bars indicate genes in the 200 Kb window surrounding SCRI_RS_235243, the red arrow indicates the DHAR (drought tolerant-related gene) hit by SCRI_RS_235243, (b) The gene structure of DHAR and the functional annotation of SNPs in this gene. (c) Haplotype structures of DHAR based on the SNPs in this gene. L: low; H: high.

## Supplemental tables

Table S1: 784 barley landraces after removing 19 accessions from the 803 Poets et al. 2015 panel.

Table S2: Details of the exome capture data from 62 landraces.

Table S3: Summary statistics for top three independent components (IC) and principal components (PCs) calculated from 19 BIOs

Table S4: The samples size for each partitions for *F*_ST_ comparisons

Table S5: Known flowering time-related genes list.

Table S6: Known cold tolerance-related genes list.

Table S7: Known drought tolerance-related genes list.

Table S8: The annotation of the 155 significant SNPs identified by environmental association. Table S9: The average and standard deviation of *F*_ST_ calculated by different comparisons Table S10: 203 *F*_ST_ outliers from elevation, high and low latitude, and growth habit comparisons. Table S11: The nine overlapping SNPs identified by *F*_ST_ outliers and association analysis approaches.

Table S12: *F*_ST_ outliers from elevation comparison in a putative inverted region reported by Fang et al. 2014 without culling SNPs in strong LD.

Table S13: Barley flowering time genes with signal of adaption, the numbers of SNPs genotyped those genes and the SNPs outlier by environmental association and *F*_ST_ outlier SNPs

## Supplemental datasets

Supplemental data 1: VCF file for the 6,152 SNPs without culling SNPs in complete LD.

Supplemental data 2: Genotype matrix with 5,800 SNPs for environmental association

Supplemental data 3: The physical positions of 9K SNPs.

Supplemental data 4: The annotations for SNPs called from exome capture resequencing data from 62 landraces.

Supplemental data 5: Phenotype matrix with 25 geographic and climatic variables for environmental association.

Supplemental data 6: Inferred ancestral status for each 9K SNP.

Supplemental data 7: Inferred ancestral status for each exome resequencing SNP from 62 landraces.

Supplemental data 8: VCF file for SNPs called from exome-capture resequencing data from 62 landraces

Supplemental data 9: All p-values and Benjamini-Hochberg FDR-values from the environmental associations for 25 variables.

Supplemental data 10: All p-values and *F*_ST_ from elevation, low and high latitude, longitude, and growth habit.

## References

Balding, D.J. (2006). A tutorial on statistical methods for population association studies. Nat Rev Genet 7: 781.

Barrero-Sicilia, C., Hernando-Amado, S., González-Melendi, P., and Carbonero, P. (2011). Structure, expression profile and subcellular localisation of four different sucrose synthase genes from barley. Planta 234: 391–403.

Beaumont, M.A., and Balding, D.J. (2004). Identifying adaptive genetic divergence among populations from genome scans. Mol Ecol 13: 969–980.

Bell, A.J., and Sejnowski, T.J. (1995). An information-maximization approach to blind separation and blind deconvolution. Neural Computation 7: 1129–1159.

Bhatia, G., Patterson, N., Sankararaman, S., and Price, A.L. (2013). Estimating and interpreting *F*ST: the impact of rare variants. Genome Res 23: 1514–1521.

Boden, S.A. et al. (2014). *EARLY FLOWERING3* regulates flowering in spring barley by mediating gibberellin production and *FLOWERING LOCUS T* expression. Plant Cell 1557–1569.

Bothmer, R.V. (1992). The wild species of *Hordeum*: relationships and potential use for improvement of cultivated barley.

Campoli, C., Shtaya, M., Davis, S.J., and von Korff, M. (2012). Expression conservation within the circadian clock of a monocot: natural variation at barley *Ppd-H1* affects circadian expression of flowering time genes, but not clock orthologs. BMC Plant Biol 12: 97.

Casao, M.C. et al. (2011). Adaptation of barley to mild winters: a role for *PPDH2*. BMC Plant Biol 11: 164.

Cavalli-Sforza, L.L. (1966). Population structure and human evolution. Proc. R. Soc. Lond. B 164: 362–379.

Chang, C.C. et al. (2015). Second-generation PLINK: rising to the challenge of larger and richer datasets. GigaScience 4: 7.

Chen, Y. et al. (2016). Overexpression of the wheat expansin gene *TaEXPA2* improved seed production and drought tolerance in transgenic tobacco plants. PloS One 11: e0153494.

Choi, D.-W., Zhu, B., and Close, T.J. (1999). The barley (*Hordeum vulgare* L.) dehydrin multigene family: sequences, allele types, chromosome assignments, and expression characteristics of 11 *Dhn* genes of cv Dicktoo. Theor Appl Genet 98: 1234–1247.

Cockram, J. et al. (2007). Haplotype analysis of vernalization loci in European barley germplasm reveals novel *VRN-H1* alleles and a predominant winter *VRN-H1/VRN-H2* multilocus haplotype. Theor Appl Genet 115: 993–1001.

Comadran, J. et al. (2012). Natural variation in a homolog of *Antirrhinum CENTRORADIALIS* contributed to spring growth habit and environmental adaptation in cultivated barley. Nat Genet 44: 1388–1392.

Dawson, I.K. et al. (2015). Barley: a translational model for adaptation to climate change. New Phytol 206: 913–931.

de Meeûs, T., and Goudet, J. (2007). A step-by-step tutorial to use HierFstat to analyse populations hierarchically structured at multiple levels. Infect Genet Evol 7: 731–735.

Durvasula, A. et al. (2016). ANGSD-wrapper: utilities for analysing next-generation sequencing data. Mol Ecol Resour 16: 1449–1454.

Eckert, A.J. et al. (2010). Patterns of population structure and environmental associations to aridity across the range of loblolly pine (*Pinus taeda* L., *Pinaceae*). Genetics 185: 969–982.

Eltayeb, A.E. et al. (2006). Enhanced tolerance to ozone and drought stresses in transgenic tobacco overexpressing dehydroascorbate reductase in cytosol. Physiol Plant 127: 57–65.

Fang, Z. et al. (2014). Two genomic regions contribute disproportionately to geographic differentiation in wild barley. G3: Genes, Genomes, Genetics 4: 1193–1203.

Fang, Z. et al. (2012). Megabase-scale inversion polymorphism in the wild ancestor of maize. Genetics 191: 883–894.

Ford, B. et al. (2016). Barley *(Hordeum vulgare)* circadian clock genes can respond rapidly to temperature in an *EARLY FLOWERING 3* – dependent manner. J Exp Bot 67: 5517–5528.

Francia, E. et al. (2004). Two loci on chromosome 5H determine low-temperature tolerance in a ‘Nure’ (winter) × ‘Tremois’ (spring) barley map. Theor Appl Genet 108: 670–680.

Gaut, B.S., Seymour, D.K., Liu, Q., and Zhou, Y. (2018). Demography and its effects on genomic variation in crop domestication. Nature Plants 4: 1.

Hansson, M., Komatsuda, T., Stein, N., and Muehlbauer, G.J. (2018). Molecular mapping and cloning of genes and QTLs. In The Barley Genome, Stein, N., and Muehlbauer, G.J., eds (New York City: Springer), pp. 139–154.

Harlan, J.R., and Zohary, D. (1966). Distribution of wild wheats and barley. Science 153: 1074–1080.

Harris, D.R., and Gosden, C. (1996). The beginnings of agriculture in western Central Asia. In The origins and spread of agriculture and pastoralism in Eurasia, Harris, D.R., ed (London: UCL Press), pp. 370–389.

Hayes, P.M. et al. (1993). Quantitative trait loci on barley (*Hordeum vulgare* L.) chromosome 7 associated with components of winterhardiness. Genome 36: 66–71.

Hijmans, R.J. et al. (2016). raster: Geographic data analysis and modeling.

Pj, H., Sr, W., Tjy, K., and Pl., M. (2018). MorrellLab/sequence_handling: Release v2.0: SNP calling with GATK 3.8.

Honsdorf, N., March, T.J., Berger, B., Tester, M., and Pillen, K. (2014). High-throughput phenotyping to detect drought tolerance QTL in wild barley introgression lines. PLoS One 9: e97047.

Hufford, M.B. et al. (2012). Comparative population genomics of maize domestication and improvement. Nat Genet 44: 808.

Consortium, I.B.G.S. (2012). A physical, genetic and functional sequence assembly of the barley genome. Nature 491: 711.

Jakob, S.S., Meister, A., and Blattner, F.R. (2004). The considerable genome size variation of *Hordeum* species (Poaceae) is linked to phylogeny, life form, ecology, and speciation rates. Mol Biol Evol 21: 860–869.

Jones, H. et al. (2008). Population-based resequencing reveals that the flowering time adaptation of cultivated barley originated east of the Fertile Crescent. Mol Biol Evol 25: 2211–2219.

Kanneganti, V., and Gupta, A.K. (2008). Overexpression of *OsiSAP8*, a member of stress associated protein (SAP) gene family of rice confers tolerance to salt, drought and cold stress in transgenic tobacco and rice. Plant Mol Biol 66: 445–462.

Kantar, M.B., Nashoba, A.R., Anderson, J.E., Blackman, B.K., and Rieseberg, L.H. (2017). The genetics and genomics of plant domestication. Bioscience 67: 971–982.

Knox, A.K. et al. (2010). *CBF* gene copy number variation at *Frost Resistance-2* is associated with levels of freezing tolerance in temperate-climate cereals. Theor Appl Genet 121: 21–35.

Konishi, S. et al. (2006). An SNP caused loss of seed shattering during rice domestication. Science 312: 1392–1396.

Kono, T.J.Y. et al. (2016). The role of deleterious substitutions in crop genomes. Mol Biol Evol 33: 2307–2317.

Kono, T.J.Y. et al. (2018). The fate of deleterious variants in a barley genomic prediction population. bioRxiv 442020.

Korneliussen, T.S., Albrechtsen, A., and Nielsen, R. (2014). ANGSD: analysis of next generation sequencing data. BMC Bioinformatics 15: 356.

Lau, O.S., and Deng, X.W. (2012). The photomorphogenic repressors *COP1* and *DET1*: 20 years later. Trends Plant Sci 17: 584–593.

Lewontin, R.C. (1988). On measures of gametic disequilibrium. Genetics 120: 849–852.

Lewontin, R.C., and Krakauer, J. (1973). Distribution of gene frequency as a test of the theory of the selective neutrality of polymorphisms. Genetics 74: 175–195.

Li, C., Zhou, A., and Sang, T. (2006). Rice domestication by reducing shattering. Science 311: 1936–1939.

Lipka, A.E. et al. (2012). GAPIT: genome association and prediction integrated tool. Bioinformatics 28: 2397–2399.

Lotterhos, K.E., and Whitlock, M.C. (2014). Evaluation of demographic history and neutral parameterization on the performance of *F*ST outlier tests. Mol Ecol 23: 2178–2192.

Lowry, D.B., and Willis, J.H. (2010). A widespread chromosomal inversion polymorphism contributes to a major life-history transition, local adaptation, and reproductive isolation. PLoS Biol 8: e1000500.

Más, P., Alabadí, D., Yanovsky, M.J., Oyama, T., and Kay, S.A. (2003). Dual role of *TOC1* in the control of circadian and photomorphogenic responses in *Arabidopsis*. Plant Cell 15: 223–236.

Mascher, M. et al. (2017). A chromosome conformation capture ordered sequence of the barley genome. Nature 544: 427–433.

Morrell, P.L., Buckler, E.S., and Ross-Ibarra, J. (2012). Crop genomics: advances and applications. Nat Rev Genet 13: 85.

Morrell, P.L., and Clegg, M.T. (2007). Genetic evidence for a second domestication of barley (*Hordeum vulgare*) east of the Fertile Crescent. Proceedings of the National Academy of Sciences 104: 3289–3294.

Morrell, P.L., Gonzales, A.M., Meyer, K.K.T., and Clegg, M.T. (2013). Resequencing data indicate a modest effect of domestication on diversity in barley: a cultigen with multiple origins. J Hered 105: 253–264.

Muñoz-Amatriaín, M. et al. (2014). The USDA barley core collection: genetic diversity, population structure, and potential for genome-wide association studies. PLoS One 9: e94688.

Muñoz-Amatriaín, M. et al. (2011). An improved consensus linkage map of barley based on flow-sorted chromosomes and single nucleotide polymorphism markers. The Plant Genome 4: 238–249.

Muñoz-Amatriaín, M. et al. (2015). Sequencing of 15 622 gene-bearing BAC s clarifies the gene-dense regions of the barley genome. The Plant Journal 84: 216–227.

Nei, M., and Maruyama, T. (1975). Letters to the editors: Lewontin-Krakauer test for neutral genes. Genetics 80: 395.

Neph, S. et al. (2012). BEDOPS: high-performance genomic feature operations. Bioinformatics 28: 1919–1920.

Nishida, H. et al. (2013). *Phytochrome C* is a key factor controlling long-day flowering in barley. Plant Physiol 163: 804–814.

Nordborg, M., and Tavaré, S. (2002). Linkage disequilibrium: what history has to tell us. Trends Genet 18: 83–90.

Osipova, S.V., Permyakov, A.V., Permyakova, M.D., Pshenichnikova, T.A., and Börner, A. (2011). Leaf dehydroascorbate reductase and catalase activity is associated with soil drought tolerance in bread wheat. Acta Physiol Plant 33: 2169–2177.

Paradis, E., Claude, J., and Strimmer, K. (2004). APE: analyses of phylogenetics and evolution in R language. Bioinformatics 20: 289–290.

Patterson, N., Price, A.L., and Reich, D. (2006). Population structure and eigenanalysis. PLoS Genetics 2: e190.

Pinhasi, R., Fort, J., and Ammerman, A.J. (2005). Tracing the origin and spread of agriculture in Europe. PLoS Biol 3: e410.

Poets, A.M. et al. (2015a). The effects of both recent and long-term selection and genetic drift are readily evident in North American barley breeding populations. G3: Genes, Genomes, Genetics 6: 609–622.

Poets, A.M., Fang, Z., Clegg, M.T., and Morrell, P.L. (2015b). Barley landraces are characterized by geographically heterogeneous genomic origins. Genome Biol 16: 173.

Pyhäjärvi, T., Hufford, M.B., Mezmouk, S., and Ross-Ibarra, J. (2013). Complex patterns of local adaptation in teosinte. Genome Biology and Evolution 5: 1594–1609.

R Core Team (2017). R: A language and environment for statistical computing. R Foundation for Statistical Computing

Reinheimer, J.L., Barr, A.R., and Eglinton, J.K. (2004). QTL mapping of chromosomal regions conferring reproductive frost tolerance in barley (*Hordeum vulgare* L.). Theor Appl Genet 109: 1267–1274.

Rellstab, C., Gugerli, F., Eckert, A.J., Hancock, A.M., and Holderegger, R. (2015). A practical guide to environmental association analysis in landscape genomics. Mol Ecol 24: 4348–4370.

Ross-Ibarra, J., Morrell, P.L., and Gaut, B.S. (2007). Plant domestication, a unique opportunity to identify the genetic basis of adaptation. Proc Natl Acad Sci USA 104: 8641–8648.

Russell, J. et al. (2016). Exome sequencing of geographically diverse barley landraces and wild relatives gives insights into environmental adaptation. Nat Genet 48: 1024.

Saghai-Maroof, M.A., Soliman, K.M., Jorgensen, R.A., and Allard, R.W.L. (1984). Ribosomal DNA spacer-length polymorphisms in barley: Mendelian inheritance, chromosomal location, and population dynamics. Proceedings of the National Academy of Sciences 81: 8014–8018.

Saisho, D., and Purugganan, M. (2007). Molecular phylogeography of domesticated barley traces expansion of agriculture in the Old World. Genetics 177: 1765–1776.

Sasaki, K., Christov, N.K., Tsuda, S., and Imai, R. (2013). Identification of a novel LEA protein involved in freezing tolerance in wheat. Plant Cell Physiol 55: 136–147.

Shin, J.-H., Blay, S., McNeney, B., and Graham, J. (2006). LDheatmap: an R function for graphical display of pairwise linkage disequilibria between single nucleotide polymorphisms. Journal of Statistical Software 16: 1–10.

Skinner, J.S. et al. (2006). Mapping of barley homologs to genes that regulate low temperature tolerance in *Arabidopsis*. Theor Appl Genet 112: 832–842.

Stephens, M., and Scheet, P. (2005). Accounting for decay of linkage disequilibrium in haplotype inference and missing-data imputation. The American Journal of Human Genetics 76: 449–462.

Stephens, M., Smith, N.J., and Donnelly, P. (2001). A new statistical method for haplotype reconstruction from population data. The American Journal of Human Genetics 68: 978–989.

Stockinger, E.J., Skinner, J.S., Gardner, K.G., Francia, E., and Pecchioni, N. (2007). Expression levels of barley *Cbf* genes at the *Frost resistance-H2* locus are dependent upon alleles at *Fr-H1* and *Fr-H2*. The Plant Journal 51: 308–321.

Sweigart, A.L., and Willis, J.H. (2003). Patterns of nucleotide diversity are affected by mating system and asymmetric introgression in two species of *Mimulus*. Evolution 57: 2490–2506.

Teshima, K.M., Coop, G., and Przeworski, M. (2006). How reliable are empirical genomic scans for selective sweeps. Genome Res 16: 702–712.

Tiffin, P., and Ross-Ibarra, J. (2014). Advances and limits of using population genetics to understand local adaptation. Trends Ecol Evol 29: 673–680.

Tondelli, A. et al. (2006). Mapping regulatory genes as candidates for cold and drought stress tolerance in barley. Theor Appl Genet 112: 445–454.

Turner, A., Beales, J., Faure, S., Dunford, R.P., and Laurie, D.A. (2005). The pseudoresponse regulator *Ppd-H1* provides adaptation to photoperiod in barley. Science 310: 1031–1034.

Visioni, A. et al. (2013). Genome-wide association mapping of frost tolerance in barley (*Hordeum vulgare* L.). BMC Genomics 14: 424.

Wagner, D.B., and Allard, R.W. (1991). Pollen migration in predominantly self-fertilizing plants: barley. J Hered 82: 302–304.

Wang, K., Li, M., and Hakonarson, H. (2010). ANNOVAR: functional annotation of genetic variants from high-throughput sequencing data. Nucleic Acids Res 38: e164–e164.

Wang, L. et al. (2017). The interplay of demography and selection during maize domestication and expansion. Genome Biol 18: 215.

Weir, B.S., and Cockerham, C.C. (1984). Estimating F-statistics for the analysis of population structure. Evolution 38: 1358–1370.

Willcox, G. (2002). Geographical variation in major cereal components and evidence for independent domestication events in western Asia. In The Dawn of Farming in the Near East, ex oriente Berlin), pp. 133–140.

Wright, S. (1949). The genetical structure of populations. Annals of Eugenics 15: 323–354.

Xu, D., Zhu, D., and Deng, X.W. (2016). The role of COP1 in repression of photoperiodic flowering. F1000Research 5:

Yan, L. et al. (2006). The wheat and barley vernalization gene *VRN3* is an orthologue of *FT*. Proc Natl Acad Sci USA 103: 19581–19586.

Yoder, J.B. et al. (2014). Genomic signature of adaptation to climate in *Medicago truncatula*. Genetics 196: 1263–1275.

Zakhrabekova, S. et al. (2012). Induced mutations in circadian clock regulator *Mat-a* facilitated short-season adaptation and range extension in cultivated barley. Proceedings of the National Academy of Sciences 109: 4326–4331.

Zhang, L. et al. (2009). Selection on grain shattering genes and rates of rice domestication. New Phytol 184: 708–720.

Zhang, Z. et al. (2010). Mixed linear model approach adapted for genome-wide association studies. Nat Genet 42: 355.

Zohary, D., Hopf, M., and Weiss, E. (2012). Domestication of plants in the Old World: the origin and spread of domesticated plants in Southwest Asia, Europe, and the Mediterranean Basin. (Oxford: Oxford University Press).

