## Supplemental notes and supplemental figures for "Environmental association identifies candidates for tolerance to low temperature and drought"

Method S1

### Physical Mapping of 9K SNPs

A number of genetic mapping studies have reported genetic map positions for barley SNPs (e.g.Comadran et al., 2012; Mayer et al., 2011; Muñoz-Amatriaín et al., 2014), and the physical locations for a portion of iSelect SNPs relative to the barley reference genome (Mascher et al., 2017) have been reported previously (Cantalapiedra et al., 2015; Colmsee et al., 2015; Comadran et al., 2012; Mayer et al., 2011). We used the contextual sequences of 7,864 SNPs from the 9K Illumina Infinium iSelect Custom Genotyping BeadChip (typically either 121 or 241 bp long) (Comadran et al., 2012) to perform BLASTn (Altschul et al., 1990) searches against the masked reference genome (Mascher et al., 2017). We configured BLASTn to reject hits with an expect value greater than 0.000001 and identity less than 90%. For sequences where BLAST did not identify a unique position, we used previously reported genetic map positions (Muñoz-Amatriaín et al., 2014) to infer the most likely chromosome of origin. If two blast hits were inferred to be within 100 kb from each other, we systematically chose the one with smaller value along the pseudomolecule as the physical position. Otherwise, we identified the physical position with the hit closest to the genetic position. The mapping of 9K SNPs was performed using the Python program SNP_Utils (https://github.com/mojaveazure/SNP_Utils). 425 SNPs were not aligned due to either no hits above the e-value threshold ≥0.000001 or identity ≤ 90% or multiple hits within ≤ 100 Kb for SNPs with no estimated genetic positions. For those 425 SNPs, we used a manual BLAST search of contextual sequence using the IPK web server (http://webblast.ipk-gatersleben.de/barley_ibsc/viroblast.php) with default parameters. The BLAST search used no threshold and involved selecting the hits with the highest combined rank of identity and score. If the contextual sequence did not have a unique BLAST hit in the genome, we used SNPMeta (Kono et al., 2014) to identify the potential genes where the SNP resides (see link to Barley_SNP_Annotations in supplemental text for SNPmeta results), then a BLAST search of the gene against the masked reference genome to identify the physical location of the best hit.

### Exome capture sequence data handling and SNP calling

Sequence quality assessment used FastQC (http://www.bioinformatics.bbsrc.ac.uk/projects/fastqc/) and seqqs (https://github.com/vsbuffalo/seqqs). Reads were counted using bioawk scripts (https://github.com/lh3/bioawk). Adapter trimming used Scythe (https://github.com/vsbuffalo/scythe). Reference-based read mapping against the draft barley reference genome (Mascher et al., 2017) was conducted with the Burrows-Wheeler Aligner (BWA-MEM) (Li and Durbin, 2009). Read mapping used default parameters for BWA-MEM except for the following: 8 threads, minimum seed length of 10, re-seed value of 1.0, a gap penalty of 8, a mismatch penalty of 4, and a minimum reporting threshold of 85. Read mapping parameters were chosen to permit a ~2% mismatch between reads and the reference sequence, the highest estimated nucleotide diversity reported based on Sanger resequencing in wild and cultivated barley (Morrell et al., 2003; Morrell et al., 2006; Morrell et al., 2013). Samtools (Li et al., 2009) and Picard (http://broadinstitute.github.io/picard/), were used for alignment sorting, de-duplicating, and adding read groups to the resulting BAM files. Estimates of read depth and coverage made use of 'bedtools genomecov' (Quinlan and Hall, 2010) relative to an empirical estimate of exome coverage (Kono et al., 2018). Briefly, this estimate was based on mapping of roughly 241X exome capture reads from Morex, reported by Mascher et al. (2017), back to the Morex draft genome (Mascher et al., 2017). Regions with > 50X were considered covered by exome capture (https://github.com/MorrellLAB/captured_50x_BED). This results in ~80 Mb of exome coverage (Kono et al., 2018) relative to the 60 Mb capture design by (Mascher et al., 2013).

Alignment processing followed the Genome Analysis Toolkit (GATK) best practices workflow (DePristo et al., 2011; McKenna et al., 2010). Cleaned BAM alignments were realigned around putative insertion/deletion (indel) sites. Individual sample genotype likelihoods were then calculated with the HaplotypeCaller, with a haploid model and a "heterozygosity" (pairwise diversity) value of 0.008 per base pair. This value is the mean estimate of coding nucleotide sequence diversity, based on previous Sanger resequencing experiments (Caldwell et al., 2006; Morrell et al., 2013). SNP calls were made from the genotype likelihoods with the GATK tool GenotypeGVCFs (McKenna et al., 2010). A high-confidence subset of these SNP calls was created by both site-based filtering (position, number of alleles, QUAL, missingness, heterozygosity) and then genotype-based filtering (genotype quality, depth). The site-based filtering was to exclude indels, sites with more than two alleles, sites with a QUAL score < 40 or missing, sites outside the exome capture regions, sites with > 90% heterozygous calls, and sites with ≥ 20% of the calls either missing. The genotype-based filtering was to filter out SNPswith a genotype quality < 9, or with a depth < 5. We used 7,864 9K SNPs with physical positions, 708,620 SNPs from (Kono et al., 2016), and 2,124,487 SNPs from wild barley as prior variants and ran GATK `Variant Recalibrator` and Apply `Recalibration` to recalibrate the variants called by GATK to produce the raw VCF file.

Sites outside the empirically inferred exome capture regions, sites with more than two alleles, and indels were filtered from the raw VCF file. Genotype calls were considered missing if the individual read depths were <5 and >109 (the 95th percentile of coverage), or genotype quality was < 9, or the ratio of reads supported for reference to alternative alleles showed a > 10% deviation from 50:50. SNP positions with > 90% heterozygous calls, a QUAL score < 40, or > 20% missing genotype calls were discarded.

Scripts to perform adapter contamination removal, read mapping, alignment cleaning, implementing the GATK best practices workflow, GATK VariantRecalibrator, ApplyRecalibration, and filtering for VCF files are part of the `sequence_handling` workflow reported by PJ et al. (2018).

### Environmental association mapping

The latitude, longitude, elevation, and BIO1 to BIO19 values of the collection locations for 784 landraces are given in the phenotype data file (Supplemental data 5). To determine if consolidation of components identifies novel variants, the top three Independent Components (IC) were calculated from BIO1 to BIO19 values after standardization of each BIO variable using the icaimax 'ica' function from the ica-package in R (Bell and Sejnowski, 1995). We summarized BIO variables with ICs rather than principal component analysis (PCA) because ICs infer the orthogonal coordinate system with a given number of dimensions that capture the data best (maximize the non-Gaussian-ness). PCA is designed to rotate the original orthogonal coordinate system around the origin to capture the highest variance of the data in the order of principal components, an approach that is highly susceptible to redundancy in the data. To compare the summaries of BIO variables, we tested for correlation between the ICs/PCs and individual Bioclim variables. We found that using the top three ICs appears to capture the cold temperature trend better than using top three PCs (Table S4). However, the ICs constitute a somewhat extreme summary of the bioclimatic variables, as the first three ICs included only eight bioclimatic variables (Table S3). Those eight variables are not closely related to the remainder of the BIO variables (Figure S5) and potentially result in a loss of information with regard to associations with remaining variables. Therefore, we report association analysis to both ICs and individual Bioclim variables (Supplemental data 5).

The GAPIT and Efficient Mixed Model Association (EMMA) packages (Lipka et al., 2012; Zhang et al., 2010) were used together with the R packages, MASS (Venables and Ripley, 2002), multtest (Pollard et al., 2004), gplots (Warnes et al., 2009), compiler (Tierney, 2019), and scatterplot3d (Ligges and Mächler, 2002), according to the GAPIT demo script (http://www.zzlab.net/GAPIT/gapit_tutorial_script.txt). For GAPIT, default parameter values were used except that the PCA total argument was set to three. We exclude SNPs with minor allele frequency (MAF) > 0.01 from association analysis. To correct for multiple testing, we applied the Benjamini-Hochberg false discovery rate (FDR) correction. We report both adjusted *p*-values and FDR with an FDR threshold ≤ 0.25.

Permutation and Empirical *P*-value for $F_{ST}$

The null distribution of $F_{ST}$ values was estimated from 1,000 permutations of genotype data for each comparison. For example, 80 randomly selected individuals were assigned to either the spring or winter partition for growth habit. Because there are only 54 samples from the high elevation > 3000 m partition, a random sample of 25 individuals was assigned to each partition for the calculation of the null $F_{ST}$ distribution. The R package Hierfstat (Goudet, J, 2005) was used to calculate $F_{ST}$ at each SNP. The *p*-value for each SNP was calculated as the percentage of null $F_{ST}$ values (out of 1,000) that exceeded the observed empirical $F_{ST}$ value.

Reference:

**Altschul, S.F., Gish, W., Miller, W., Myers, E.W., and Lipman, D.J.** (1990). Basic local alignment search tool. J Mol Biol **215**: 403–410.

**Bell, A.J., and Sejnowski, T.J.** (1995). An information-maximization approach to blind separation and blind deconvolution. Neural Computation **7**: 1129–1159.

**Caldwell, K.S., Russell, J., Langridge, P., and Powell, W.** (2006). Extreme population-dependent linkage disequilibrium detected in an inbreeding plant species, Hordeum vulgare. Genetics **172**: 557–567.

**Cantalapiedra, C.P., Boudiar, R., Casas, A.M., Igartua, E., and Contreras-Moreira, B.** (2015). BARLEYMAP: physical and genetic mapping of nucleotide sequences and annotation of surrounding loci in barley. Mol Breed **35**: 13.

**Colmsee, C., Beier, S., Himmelbach, A., Schmutzer, T., Stein, N., Scholz, U., and Mascher, M.** (2015). BARLEX–the barley draft genome explorer. Molecular plant **8**: 964–966.

**Comadran, J. et al.** (2012). Natural variation in a homolog of *Antirrhinum CENTRORADIALIS* contributed to spring growth habit and environmental adaptation in cultivated barley. Nat Genet **44**: 1388–1392.

**DePristo, M.A., Banks, E., Poplin, R., Garimella, K.V., Maguire, J.R., Hartl, C., Philippakis, A.A., Del Angel, G., Rivas, M.A., and Hanna, M.** (2011). A framework for variation discovery and genotyping using next-generation DNA sequencing data. Nat Genet **43**: 491.

**PJ, H., SR, W., TJY, K., and PL., M.** (2018). MorrellLab/sequence_handling: Release v2.0: SNP calling with GATK 3.8.

**Kono, T.J.Y., Fu, F., Mohammadi, M., Hoffman, P.J., Liu, C., Stupar, R.M., Smith, K.P., Tiffin, P., Fay, J.C., and Morrell, P.L.** (2016). The role of deleterious substitutions in crop genomes. Mol Biol Evol **33**: 2307–2317.

**Kono, T.J.Y., Liu, C., Vonderharr, E.E., Koenig, D., Fay, J.C., Smith, K.P., and Morrell, P.L.** (2018). The fate of deleterious variants in a barley genomic prediction population. bioRxiv 442020.

**Kono, T.J.Y., Seth, K., Poland, J.A., and Morrell, P.L.** (2014). SNPM eta: SNP annotation and SNP metadata collection without a reference genome. Mol Ecol Resour **14**: 419–425.

**Li, H., and Durbin, R.** (2009). Fast and accurate short read alignment with Burrows–Wheeler transform. bioinformatics **25**: 1754–1760.

**Li, H., Handsaker, B., Wysoker, A., Fennell, T., Ruan, J., Homer, N., Marth, G., Abecasis, G., and Durbin, R.** (2009). The sequence alignment/map format and SAMtools. Bioinformatics **25**: 2078–2079.

**Ligges, U., and Mächler, M.** (2002). Scatterplot3d-an r package for visualizing multivariate data.

**Lipka, A.E., Tian, F., Wang, Q., Peiffer, J., Li, M., Bradbury, P.J., Gore, M.A., Buckler, E.S., and Zhang, Z.** (2012). GAPIT: genome association and prediction integrated tool. Bioinformatics **28**: 2397–2399.

**Mascher, M. et al.** (2017). A chromosome conformation capture ordered sequence of the barley genome. Nature **544**: 427–433.

**Mascher, M., Richmond, T.A., Gerhardt, D.J., Himmelbach, A., Clissold, L., Sampath, D., Ayling, S., Steuernagel, B., Pfeifer, M., and D'ascenzo, M.** (2013). Barley whole exome capture: a tool for genomic research in the genus Hordeum and beyond. The Plant Journal **76**: 494–505.

**Mayer, K.F.X., Martis, M., Hedley, P.E., Šimková, H., Liu, H., Morris, J.A., Steuernagel, B., Taudien, S., Roessner, S., and Gundlach, H.** (2011). Unlocking the barley genome by chromosomal and comparative genomics. The Plant Cell tpc. 110.082537.

**McKenna, A., Hanna, M., Banks, E., Sivachenko, A., Cibulskis, K., Kernytsky, A., Garimella, K., Altshuler, D., Gabriel, S., and Daly, M.** (2010). The Genome Analysis Toolkit: a MapReduce framework for analyzing next-generation DNA sequencing data. Genome Res

**Morrell, P.L., Gonzales, A.M., Meyer, K.K.T., and Clegg, M.T.** (2013). Resequencing data indicate a modest effect of domestication on diversity in barley: a cultigen with multiple origins. J Hered **105**: 253–264.

**Morrell, P.L., Lundy, K.E., and Clegg, M.T.** (2003). Distinct geographic patterns of genetic diversity are maintained in wild barley (Hordeum vulgare ssp. spontaneum) despite migration. Proceedings of the National Academy of Sciences **100**: 10812–10817.

**Morrell, P.L., Toleno, D.M., Lundy, K.E., and Clegg, M.T.** (2006). Estimating the contribution of mutation, recombination, and gene conversion in the generation of haplotypic diversity. Genetics

**Muñoz-Amatriaín, M., Cuesta-Marcos, A., Endelman, J.B., Comadran, J., Bonman, J.M., Bockelman, H.E., Chao, S., Russell, J., Waugh, R., Hayes, P.M., and Muehlbauer, G.J.** (2014). The USDA barley core collection: genetic diversity, population structure, and potential for genome-wide association studies. PLoS One **9**: e94688.

**Pollard, K.S., Dudoit, S., and van der Laan, M.J.** (2004). Multiple testing procedures: R multtest package and applications to genomics.

**Quinlan, A.R., and Hall, I.M.** (2010). BEDTools: a flexible suite of utilities for comparing genomic features. Bioinformatics **26**: 841–842.

**Tierney, L.** (2019). A byte code compiler for R. system **6**: 0.010.

**Venables, W.N., and Ripley, B.D.** (2002). Tree-based methods. In Modern Applied Statistics with S, Springer), pp. 251–269.

**Warnes, G.R., Bolker, B., Bonebakker, L., Gentleman, R., Huber, W., Liaw, A., Lumley, T., Maechler, M., Magnusson, A., and Moeller, S.** (2009). gplots: Various R programming tools for plotting data. R package version **2**: 1.

**Zhang, Z., Ersoz, E., Lai, C.-Q., Todhunter, R.J., Tiwari, H.K., Gore, M.A., Bradbury, P.J., Yu, J., Arnett, D.K., and Ordovas, J.M.** (2010). Mixed linear model approach adapted for genome-wide association studies. Nat Genet **42**: 355.

Supplemental figures

Figure S1: The distribution of the pairwise genetic distance (Manhattan distance) from 784 barley landraces.

Figure S2: Exome capture target density (dark blue line), crossover rate in cM/Mb (purple line), the genomic distribution of SNPs identified in 62 barley landraces (vertical light blue lines), and 9K iSelect SNPs (red triangle) for seven chromosomes. Crossover rates were calculated using 9K SNPs. SNP genetic positions are based on the genetic map of Muñoz-Amatriaín et al. (2011).

Figure S3: Relationship of barley landrace accessions based on principal components. PC1 and PC2 are depicted relative to a map of the landrace distribution in Africa and Eurasia.

Figure S4: The derived site frequency spectrum (SNPs with inferred ancestral state) for: (a) 2,806 SNPs from the 9K iSelect genotyped in the 784 landraces and (b) 340,260 SNPs with exome-capture resequencing data in 62 landraces. Ancestral state was based on majority state from *H. murinum spp. glaucum* resequencing mapped to the Morex assembly. For all SNPs: (c) Minor allele frequency, for 6,152 SNPs from the 9K iSelect and (d) for 482,714 SNPs from exome-capture resequencing.

Figure S5: The heat map of pairwise correlation coefficient (Pearson correlation) of 22 environmental variables.

Figure S6: The genomic distribution of outlier SNPs identified according to the $F_{ST}$ comparisons of elevation (below 3,000 m versus above 3,000 m), low latitude (below 30˚N versus 30-40˚N), high latitude (30-40˚N vs above 40˚N), growth habit (winter versus spring) and association analysis of 21 bioclimatic variables, which are categorized into three classes (precipitation, temperature, and geographic variables) and ICs.

Figure S7: The distribution of ranked $F_{ST}$ from two- and three-level comparisons of elevation and latitude.

Figure S8: Venn diagram for the $F_{ST}$ outliers from the comparisons of elevation, high and low latitude, and growth habit.

Figure S9: The geographic distribution of the SNPs with high $F_{ST}$. (a) The geographic distribution of allelic types of 9K SNP SCRI_RS_153793 with highest $F_{ST}$ = 0.505. The $F_{ST}$ was from the low latitude (LL) comparison. (b) The geographic distribution of allelic types of 9K SCRI_RS_134850 with highest $F_{ST}$ = 0.390. The $F_{ST}$ was from growth habit (GH) comparison. The color bar indicates the elevations in meters. The filled pink circles indicate the derived allele, while the blue open circles indicate the ancestral allele.

Figure S10: Venn diagram for the candidate SNPs significant associated with three categories of environmental variables: precipitation, temperature, and geographic factors.

Figure S11: The difference between the replaced SNPs and queried SNPs not in the exome-capture data. (a) The minor allele frequency (MAF); (b) The physical distance. The replaced

SNPs refer to those SNPs called based on the exome capture resequnecing data and with the similar MAF as the queried SNPs. But the queried SNPs are not in the exome capture resequnecing data.

Figure S12: LD decay plot for 200 Kb window around the significant SNPs associated with environmental variables. The blue bars underneath the x-axis are the annotated genes in the 200 Kb windows. The vertical dashed lines are candidate SNPlocations. The negative signs on the x-axis refer to positions downstream of the candidate SNP.

Figure S13: (a) The linkage disequilibrium (LD) analysis of candidate SNP SCRI_RS_137464 significant associated with ”min temperature of coldest month (BIO6)” and ”mean temperature of the coldest quarter (BIO6 and 11)”. The blue bars indicate genes in the 200 Kb window surrounding 11_10380, the red arrow indicates the WCI 16 (cold tolerance-related gene) hit by 11_10380, (b) The gene structure of *WCI 16* and the functional annotation of SNPs in this gene. (c) Haplotype structures of *WCI 16* based on the SNPs in this gene. L: low; H: high.

Figure S14: (a) The linkage disequilibrium (LD) analysis of candidate SNP SCRI_RS_235243 significantly associated with ”precipitation of driest months” (BIO14). The blue bars indicate genes in the 200 Kb window surrounding SCRI_RS_235243, the red arrow indicates the DHAR (drought tolerant-related gene) hit by SCRI_RS_235243, (b) The gene structure of DHAR and the functional annotation of SNPs in this gene. (c) Haplotype structures of DHAR based on the SNPs in this gene. L: low; H: high.

Supplemental tables:

Table S1: 784 barley landraces after removing 19 accessions from the 803 Poets et al. 2015 panel.
Table S2: Details of the exome capture data from 62 landraces.
Table S3: Summary statistics for top three independent components (IC) and principal components (PCs) calculated from 19 BIOs
Table S4: The samples size for each partitions for $F_{ST}$ comparisons

Table S5: Known flowering time-related genes list.

Table S6: Known cold tolerance-related genes list.

Table S7: Known drought tolerance-related genes list.

Table S8: The annotation of the 155 significant SNPs identified by environmental association.

Table S9: The average and standard deviation of $F_{ST}$ calculated by different comparisons

Table S10: 203 $F_{ST}$ outliers from elevation, high and low latitude, and growth habit comparisons.

Table S11: The nine overlapping SNPs identified by $F_{ST}$ outliers and association analysis approaches.

Table S12: $F_{ST}$ outliers from elevation comparison in a putative inverted region reported by Fang et al. 2014 without culling SNPs in strong LD.

Table S13: Barley flowering time genes with signal of adaption, the numbers of SNPs genotyped those genes and the SNPs outlier by environmental association and $F_{ST}$ outlier SNPs

Supplemental datasets:

Supplemental data 1: VCF file for the 6,152 SNPs without culling SNPs in complete LD.

Supplemental data 2: Genotype matrix with 5,800 SNPs for environmental association

Supplemental data 3: The physical positions of 9K SNPs.

Supplemental data 4: The annotations for SNPs called from exome capture resequencing data from 62 landraces.

Supplemental data 5: Phenotype matrix with 25 geographic and climatic variables for environmental association.

Supplemental data 6: Inferred ancestral status for each 9K SNP.

Supplemental data 7: Inferred ancestral status for each exome resequencing SNP from 62 landraces.

Supplemental data 8: VCF file for SNPs called from exome-capture resequencing data from 62 landraces

Supplemental data 9: All *p*-values and Benjamini-Hochberg FDR-values from the environmental associations for 25 variables.

Supplemental data 10: All *p*-values and $F_{ST}$ from elevation, low and high latitude, longitude, and growth habit.

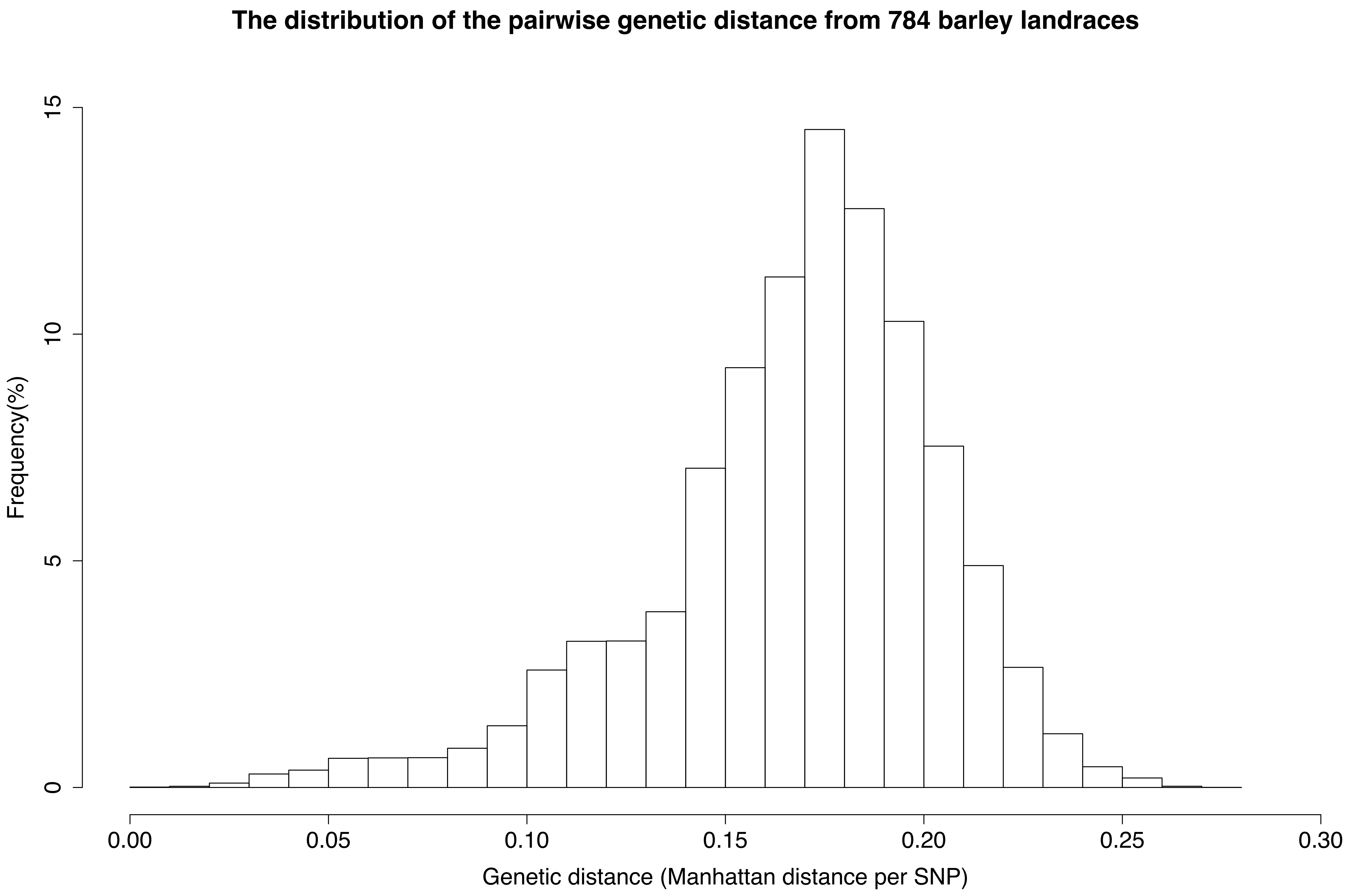

The distribution of the pairwise genetic distance from 784 barley landraces
Frequency(%)
0
5
10
15
0.00
0.05
0.10
0.15
0.20
0.25
0.30
Genetic distance (Manhattan distance per SNP)


**Figure S1**

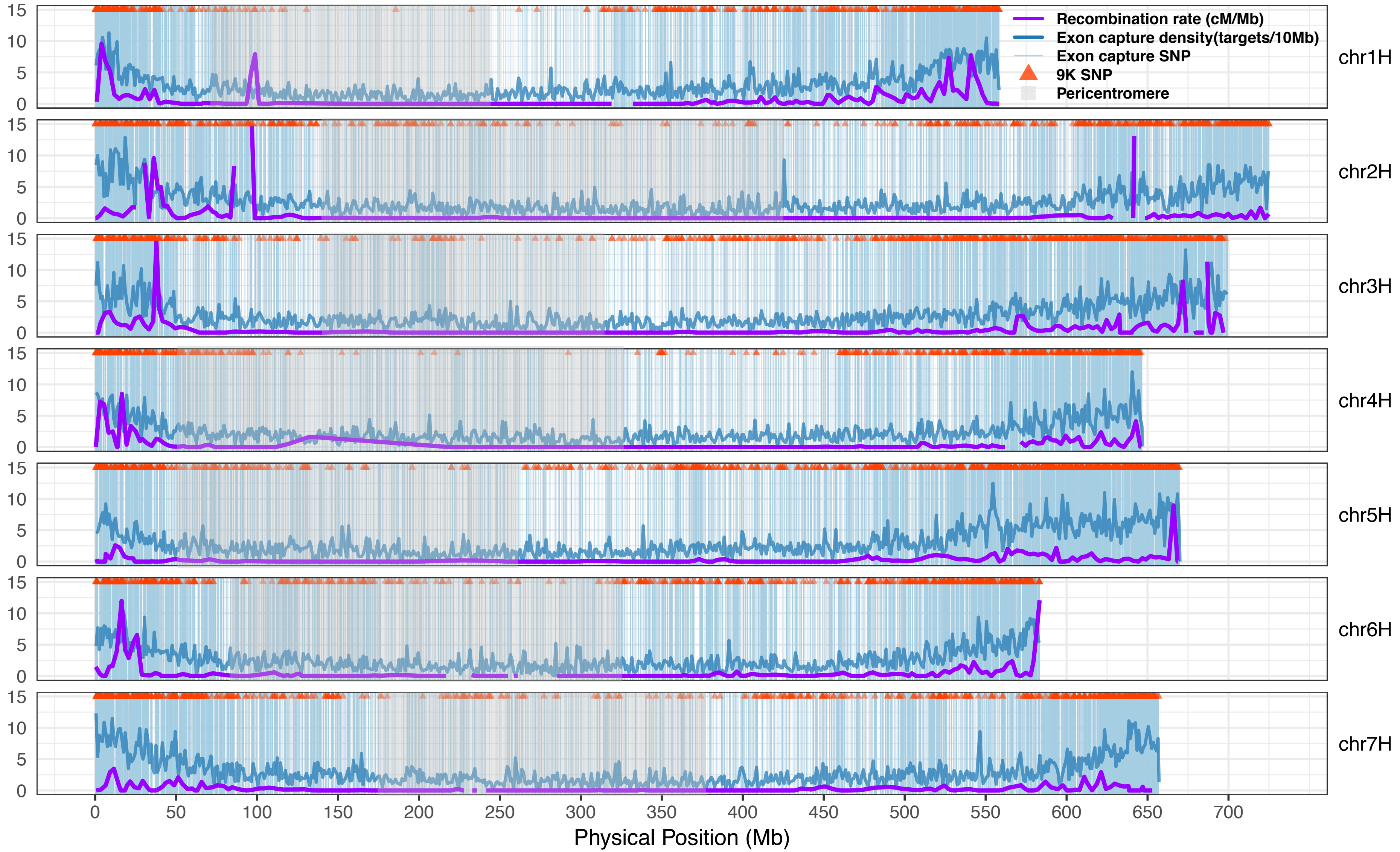

Recombination rate (cM/Mb)
Exon capture density(targets/10Mb)
Exon capture SNP
9K SNP
Pericentromere
chr1H
chr2H
chr3H
chr4H
chr5H
chr6H
chr7H
15
10
5
0
0
50
100
150
200
250
300
350
400
450
500
550
600
650
700
Physical Position (Mb)


**Figure S2**

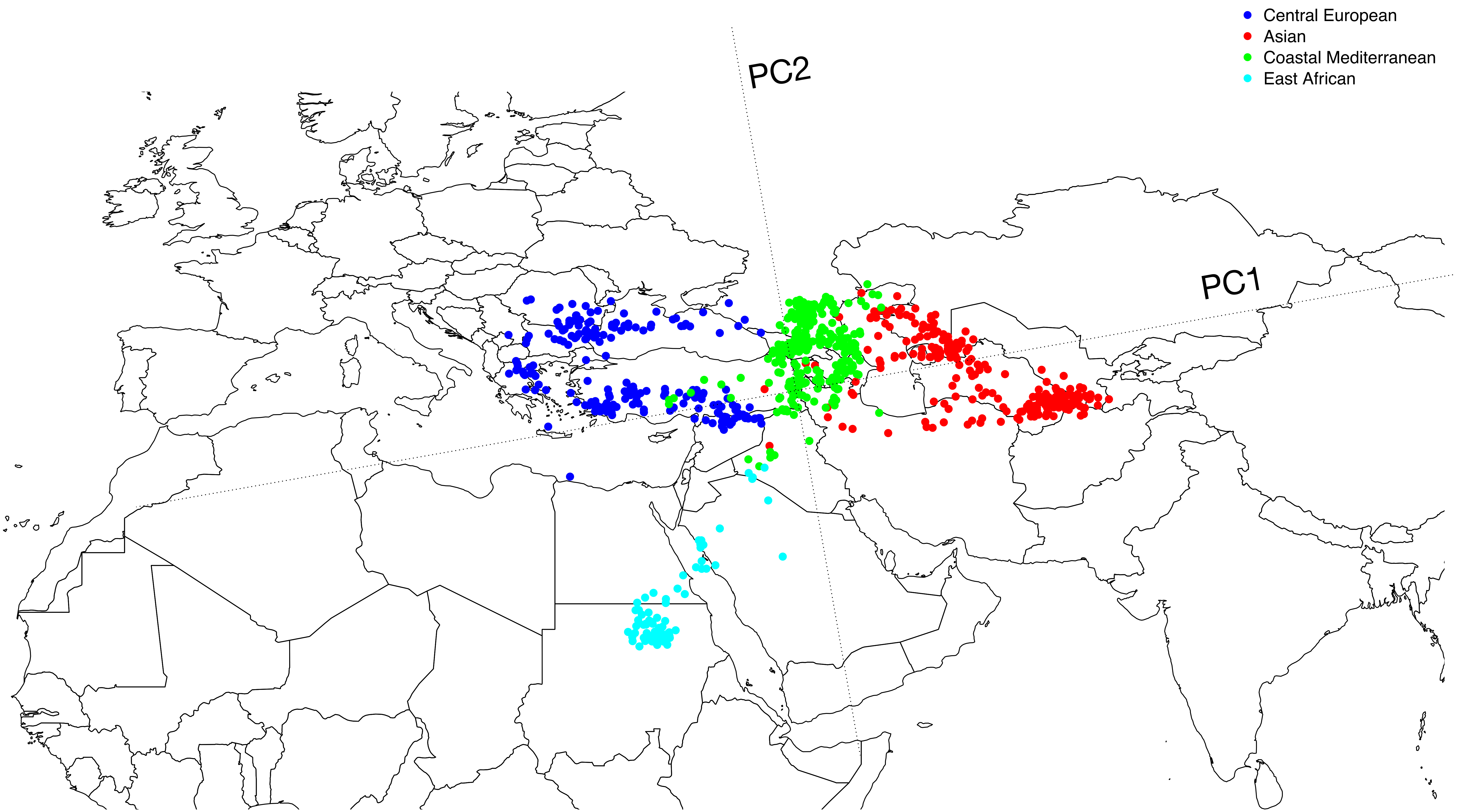
Central European
Asian
Coastal Mediterranean
East African
PC2
PC1

**Figure S3**

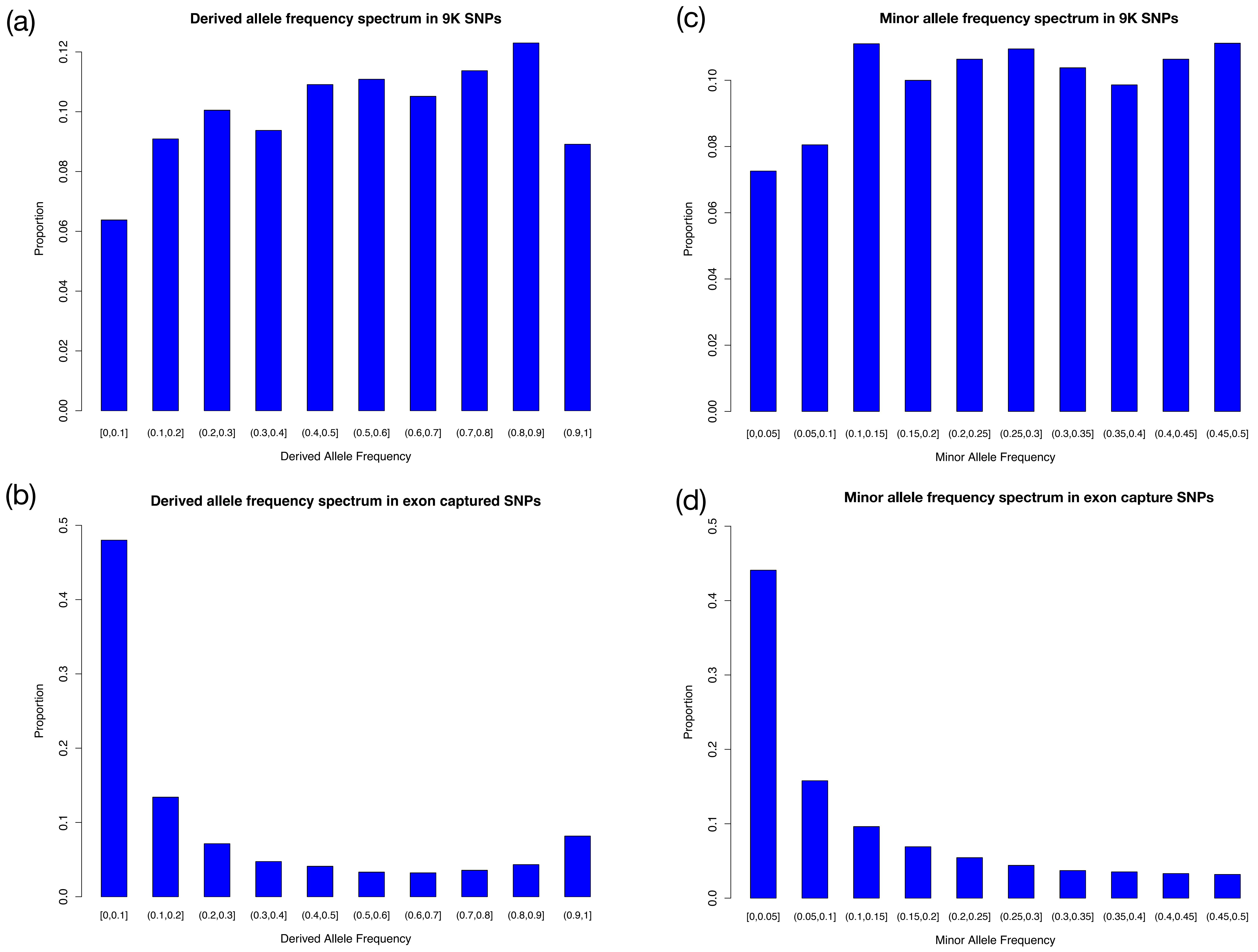
(a)
Derived allele frequency spectrum in 9K SNPs
Proportion
0.00
0.02
0.04
0.06
0.08
0.10
0.12
[0,0.1]
(0.1,0.2]
(0.2,0.3]
(0.3,0.4]
(0.4,0.5]
(0.5,0.6]
(0.6,0.7]
(0.7,0.8]
(0.8,0.9]
(0.9,1]
Derived Allele Frequency
(c)
Minor allele frequency spectrum in 9K SNPs
Proportion
0.00
0.02
0.04
0.06
0.08
0.10
[0,0.05]
(0.05,0.1]
(0.1,0.15]
(0.15,0.2]
(0.2,0.25]
(0.25,0.3]
(0.3,0.35]
(0.35,0.4]
(0.4,0.45]
(0.45,0.5]
Minor Allele Frequency
(b)
Derived allele frequency spectrum in exon captured SNPs
Proportion
0.0
0.1
0.2
0.3
0.4
0.5
[0,0.1]
(0.1,0.2]
(0.2,0.3]
(0.3,0.4]
(0.4,0.5]
(0.5,0.6]
(0.6,0.7]
(0.7,0.8]
(0.8,0.9]
(0.9,1]
Derived Allele Frequency
(d)
Minor allele frequency spectrum in exon capture SNPs
Proportion
0.0
0.1
0.2
0.3
0.4
0.5
[0,0.05]
(0.05,0.1]
(0.1,0.15]
(0.15,0.2]
(0.2,0.25]
(0.25,0.3]
(0.3,0.35]
(0.35,0.4]
(0.4,0.45]
(0.45,0.5]
Minor Allele Frequency

**Figure S4**

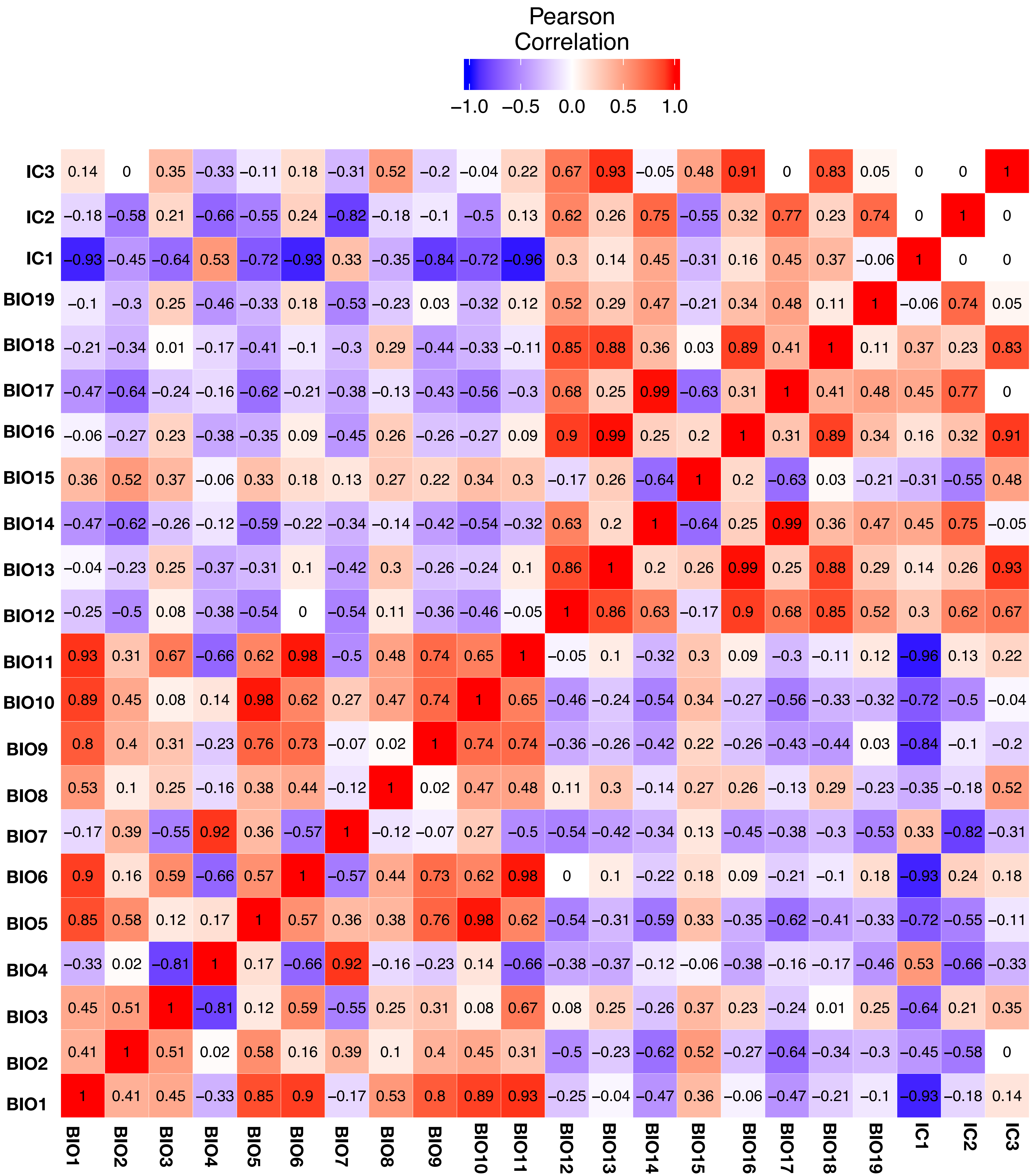
Pearson Correlation
−1.0 −0.5 0.0 0.5 1.0
IC3 0.14 0 0.35 −0.33 −0.11 0.18 −0.31 0.52 −0.2 −0.04 0.22 0.67 0.93 −0.05 0.48 0.91 0 0.83 0.05 0 0 1
IC2 −0.18 −0.58 0.21 −0.66 −0.55 0.24 −0.82 −0.18 −0.1 −0.5 0.13 0.62 0.26 0.75 −0.55 0.32 0.77 0.23 0.74 0 1 0
IC1 −0.93 −0.45 −0.64 0.53 −0.72 −0.93 0.33 −0.35 −0.84 −0.72 −0.96 0.3 0.14 0.45 −0.31 0.16 0.45 0.37 −0.06 1 0 0
BIO19 −0.1 −0.3 0.25 −0.46 −0.33 0.18 −0.53 −0.23 0.03 −0.32 0.12 0.52 0.29 0.47 −0.21 0.34 0.48 0.11 1 −0.06 0.74 0.05
BIO18 −0.21 −0.34 0.01 −0.17 −0.41 −0.1 −0.3 0.29 −0.44 −0.33 −0.11 0.85 0.88 0.36 0.03 0.89 0.41 1 0.11 0.37 0.23 0.83
BIO17 −0.47 −0.64 −0.24 −0.16 −0.62 −0.21 −0.38 −0.13 −0.43 −0.56 −0.3 0.68 0.25 0.99 −0.63 0.31 1 0.41 0.48 0.45 0.77 0
BIO16 −0.06 −0.27 0.23 −0.38 −0.35 0.09 −0.45 0.26 −0.26 −0.27 0.09 0.9 0.99 0.25 0.2 1 0.31 0.89 0.34 0.16 0.32 0.91
BIO15 0.36 0.52 0.37 −0.06 0.33 0.18 0.13 0.27 0.22 0.34 0.3 −0.17 0.26 −0.64 1 0.2 −0.63 0.03 −0.21 −0.31 −0.55 0.48
BIO14 −0.47 −0.62 −0.26 −0.12 −0.59 −0.22 −0.34 −0.14 −0.42 −0.54 −0.32 0.63 0.2 1 −0.64 0.25 0.99 0.36 0.47 0.45 0.75 −0.05
BIO13 −0.04 −0.23 0.25 −0.37 −0.31 0.1 −0.42 0.3 −0.26 −0.24 0.1 0.86 1 0.2 0.26 0.99 0.25 0.88 0.29 0.14 0.26 0.93
BIO12 −0.25 −0.5 0.08 −0.38 −0.54 0 −0.54 0.11 −0.36 −0.46 −0.05 1 0.86 0.63 −0.17 0.9 0.68 0.85 0.52 0.3 0.62 0.67
BIO11 0.93 0.31 0.67 −0.66 0.62 0.98 −0.5 0.48 0.74 0.65 1 −0.05 0.1 −0.32 0.3 0.09 −0.3 −0.11 0.12 −0.96 0.13 0.22
BIO10 0.89 0.45 0.08 0.14 0.98 0.62 0.27 0.47 0.74 1 0.65 −0.46 −0.24 −0.54 0.34 −0.27 −0.56 −0.33 −0.32 −0.72 −0.5 −0.04
BIO9 0.8 0.4 0.31 −0.23 0.76 0.73 −0.07 0.02 1 0.74 0.74 −0.36 −0.26 −0.42 0.22 −0.26 −0.43 −0.44 0.03 −0.84 −0.1 −0.2
BIO8 0.53 0.1 0.25 −0.16 0.38 0.44 −0.12 1 0.02 0.47 0.48 0.11 0.3 −0.14 0.27 0.26 −0.13 0.29 −0.23 −0.35 −0.18 0.52
BIO7 −0.17 0.39 −0.55 0.92 0.36 −0.57 1 −0.12 −0.07 0.27 −0.5 −0.54 −0.42 −0.34 0.13 −0.45 −0.38 −0.3 −0.53 0.33 −0.82 −0.31
BIO6 0.9 0.16 0.59 −0.66 0.57 1 −0.57 0.44 0.73 0.62 0.98 0 0.1 −0.22 0.18 0.09 −0.21 −0.1 0.18 −0.93 0.24 0.18
BIO5 0.85 0.58 0.12 0.17 1 0.57 0.36 0.38 0.76 0.98 0.62 −0.54 −0.31 −0.59 0.33 −0.35 −0.62 −0.41 −0.33 −0.72 −0.55 −0.11
BIO4 −0.33 0.02 −0.81 1 0.17 −0.66 0.92 −0.16 −0.23 0.14 −0.66 −0.38 −0.37 −0.12 −0.06 −0.38 −0.16 −0.17 −0.46 0.53 −0.66 −0.33
BIO3 0.45 0.51 1 −0.81 0.12 0.59 −0.55 0.25 0.31 0.08 0.67 0.08 0.25 −0.26 0.37 0.23 −0.24 0.01 0.25 −0.64 0.21 0.35
BIO2 0.41 1 0.51 0.02 0.58 0.16 0.39 0.1 0.4 0.45 0.31 −0.5 −0.23 −0.62 0.52 −0.27 −0.64 −0.34 −0.3 −0.45 −0.58 0
BIO1 1 0.41 0.45 −0.33 0.85 0.9 −0.17 0.53 0.8 0.89 0.93 −0.25 −0.04 −0.47 0.36 −0.06 −0.47 −0.21 −0.1 −0.93 −0.18 0.14
BIO1 BIO2 BIO3 BIO4 BIO5 BIO6 BIO7 BIO8 BIO9 BIO10 BIO11 BIO12 BIO13 BIO14 BIO15 BIO16 BIO17 BIO18 BIO19 IC1 IC2 IC3

**Figure S5**

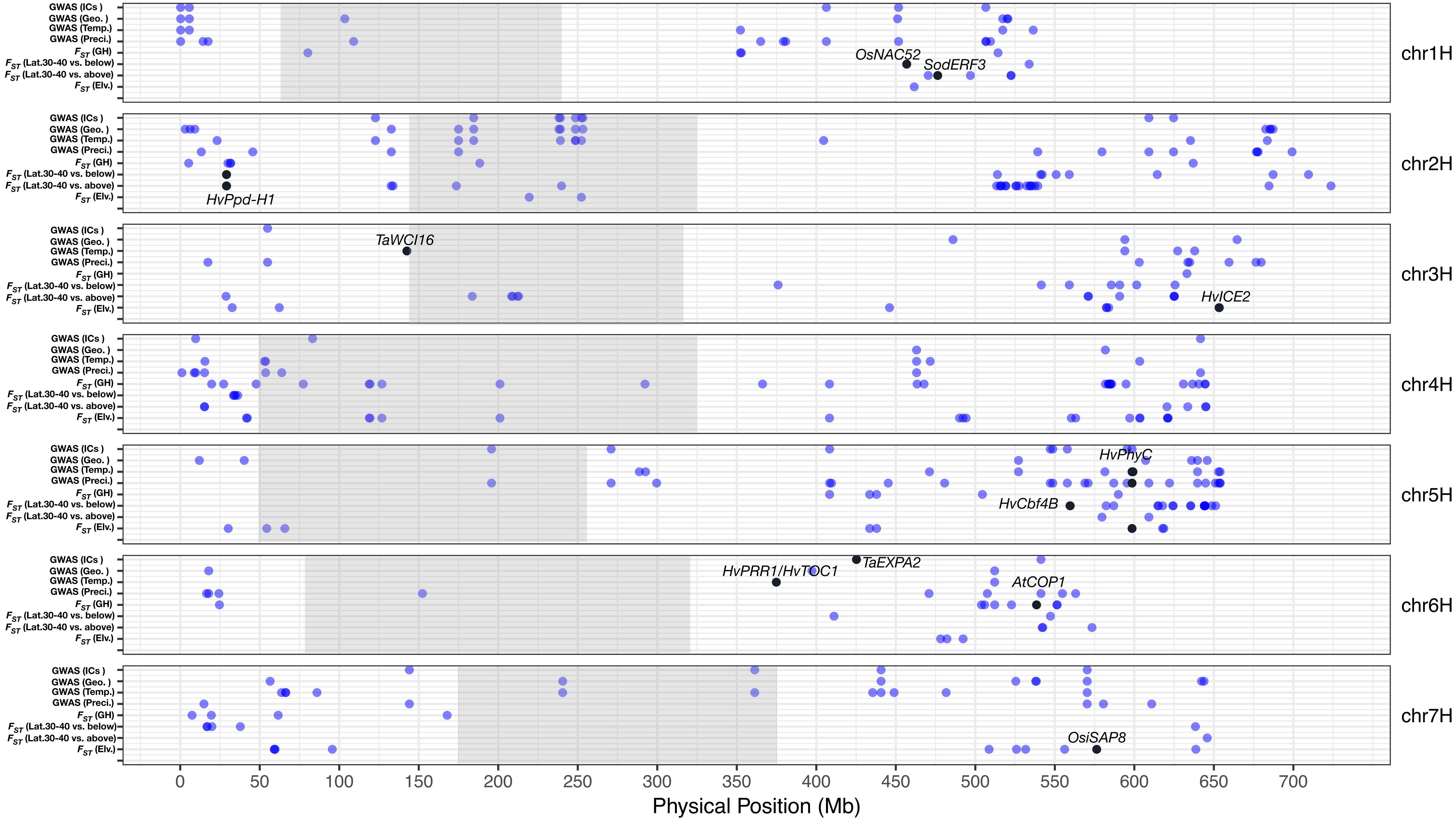

GWAS (ICs )
GWAS (Geo. )
GWAS (Temp.)
GWAS (Preci.)
$F_{ST}$ (GH)
$F_{ST}$ (Lat.30-40 vs. below)
$F_{ST}$ (Lat.30-40 vs. above)
$F_{ST}$ (Elv.)
chr1H
chr2H
chr3H
chr4H
chr5H
chr6H
chr7H
OsNAC52
SodERF3
HvPpd-H1
TaWCI16
HvICE2
HvPhyC
HvCbf4B
TaEXPA2
HvPRR1/HvTOC1
AtCOP1
OsiSAP8
0
50
100
150
200
250
300
350
400
450
500
550
600
650
700
Physical Position (Mb)


**Figure S6**

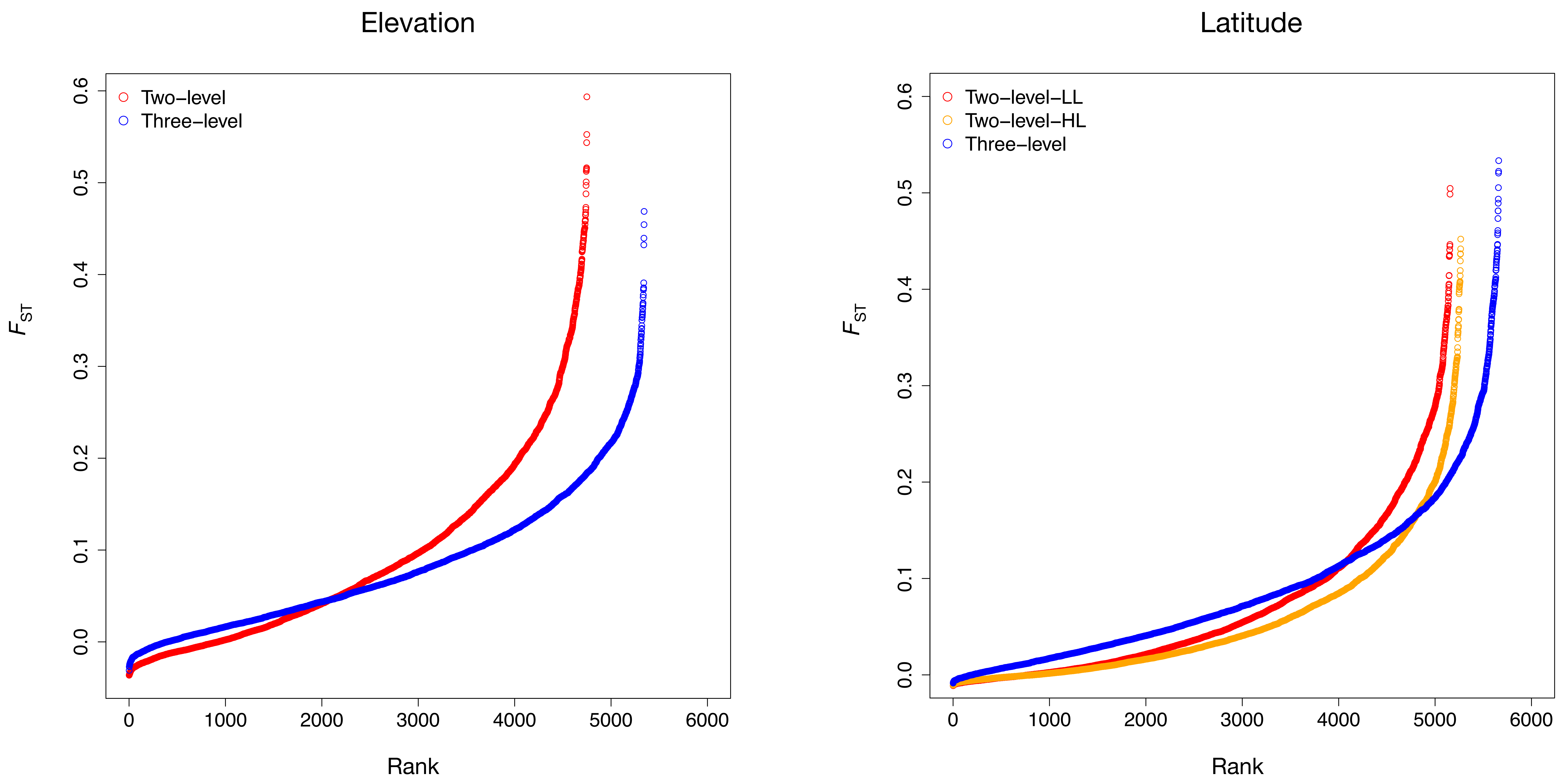
Elevation
Two–level
Three–level
Latitude
Two–level–LL
Two–level–HL
Three–level
FST
Rank
0.0
0.1
0.2
0.3
0.4
0.5
0.6
0
1000
2000
3000
4000
5000
6000

**Figure S7**

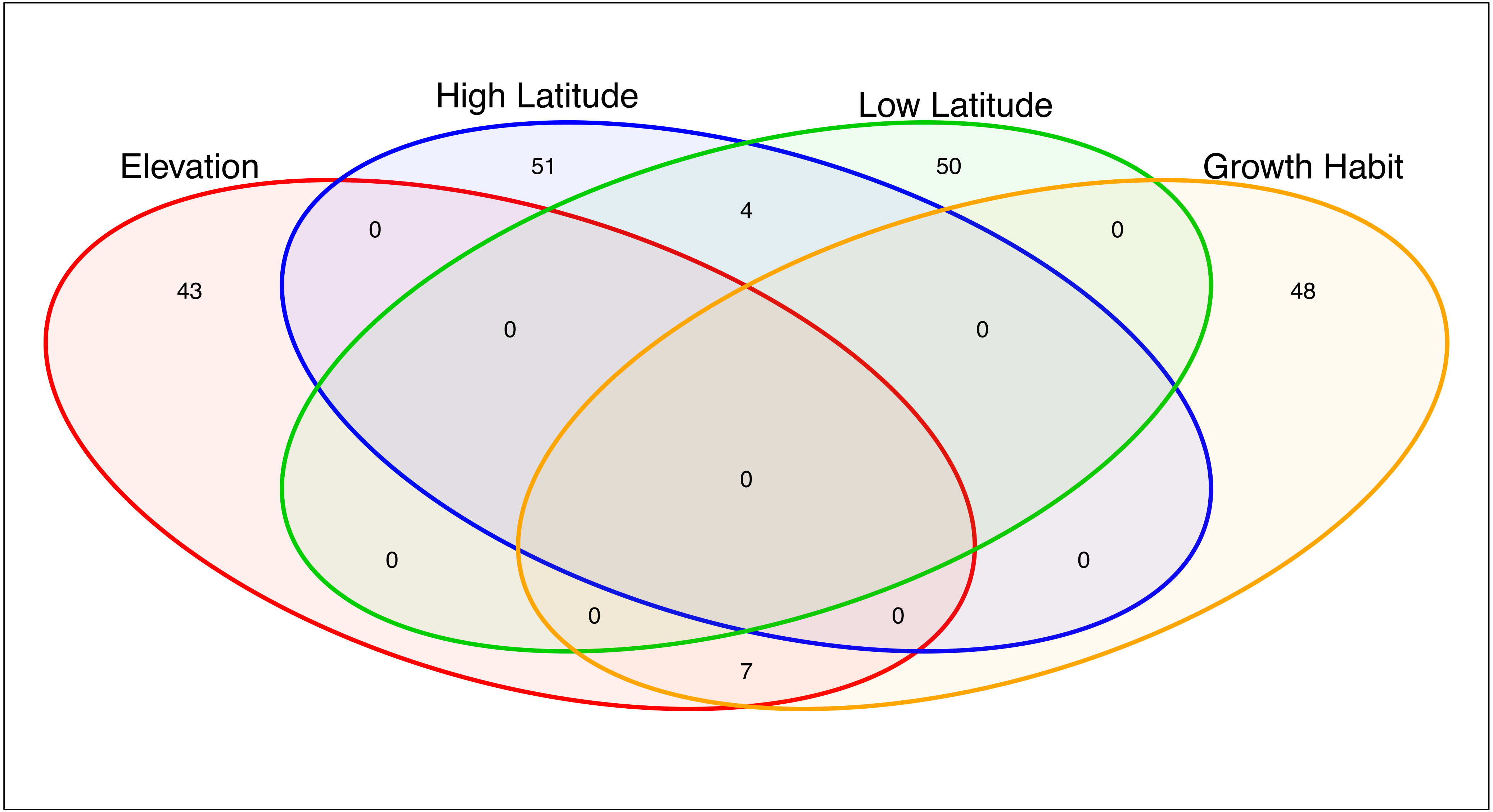
High Latitude
Low Latitude
Elevation
Growth Habit
51
50
0
4
0
43
48
0
0
0
0
0
0
0
7

**Figure S8**

(a)

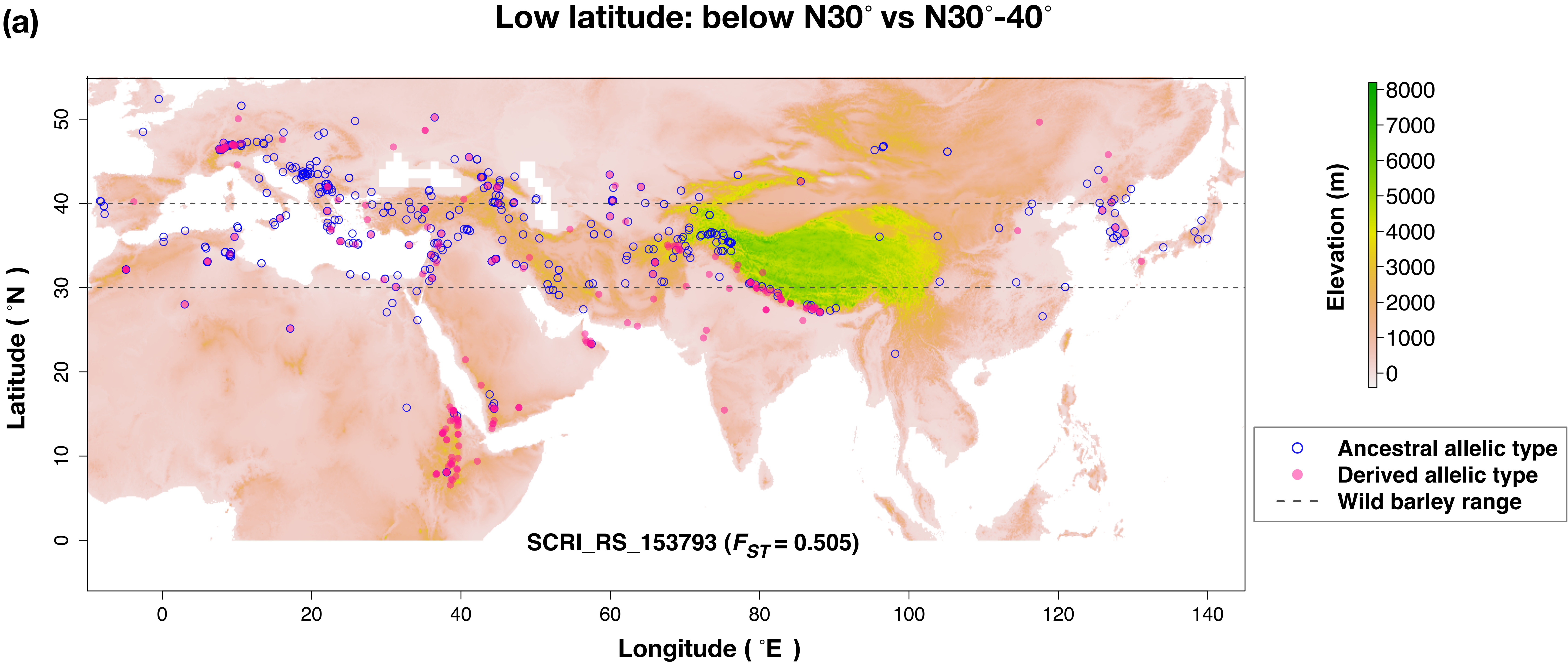
Low latitude: below N30° vs N30°-40°
Latitude ( °N )
Longitude ( °E )
50
40
30
20
10
0
0
20
40
60
80
100
120
140
SCRI_RS_153793 ($F_{ST}$ = 0.505)
Elevation (m)
8000
7000
6000
5000
4000
3000
2000
1000
0
Ancestral allelic type
Derived allelic type
Wild barley range

(b)

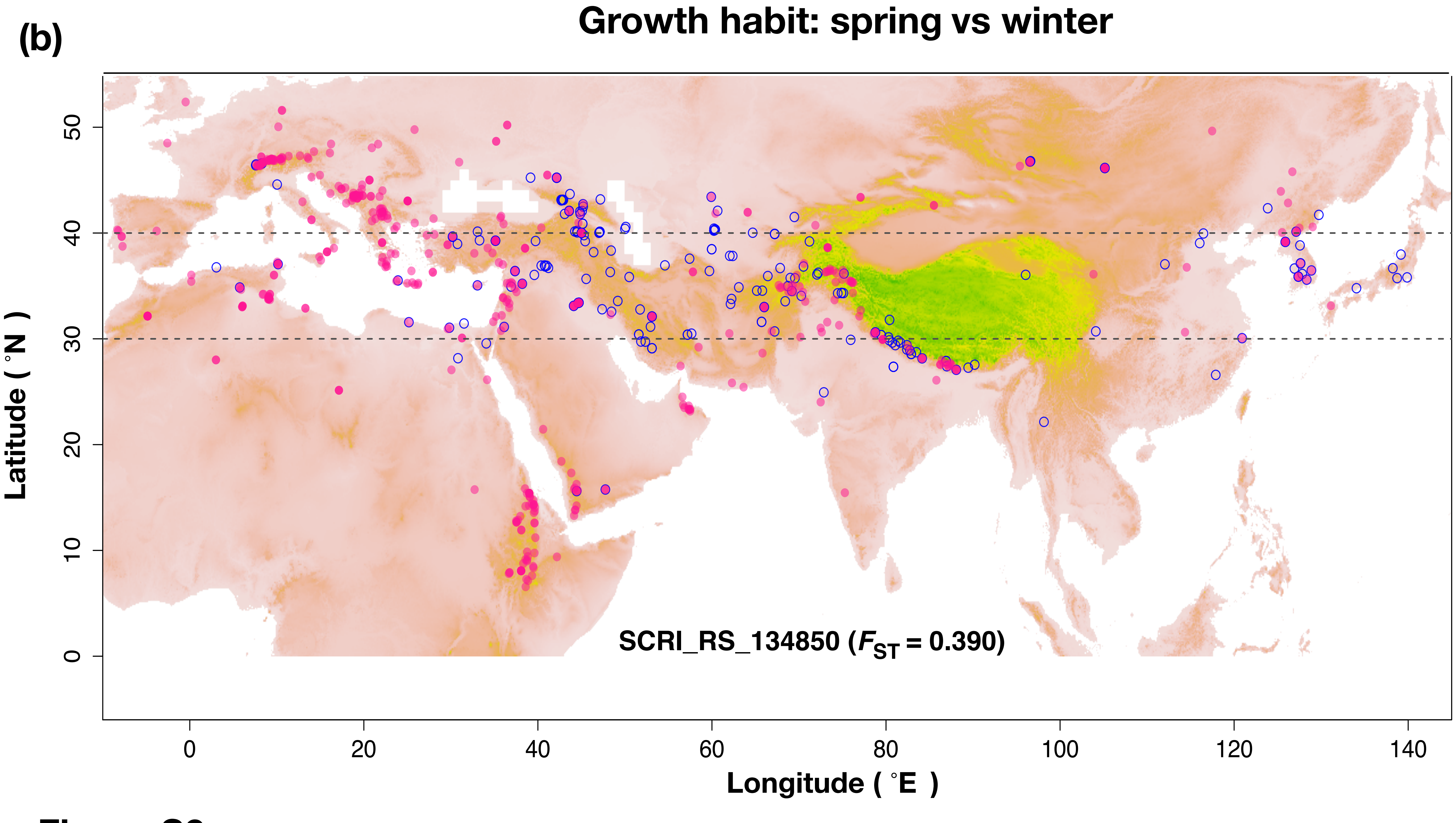
Growth habit: spring vs winter
Latitude ( °N )
Longitude ( °E )
50
40
30
20
10
0
0
20
40
60
80
100
120
140
SCRI_RS_134850 ($F_{ST}$ = 0.390)

**Figure S9**

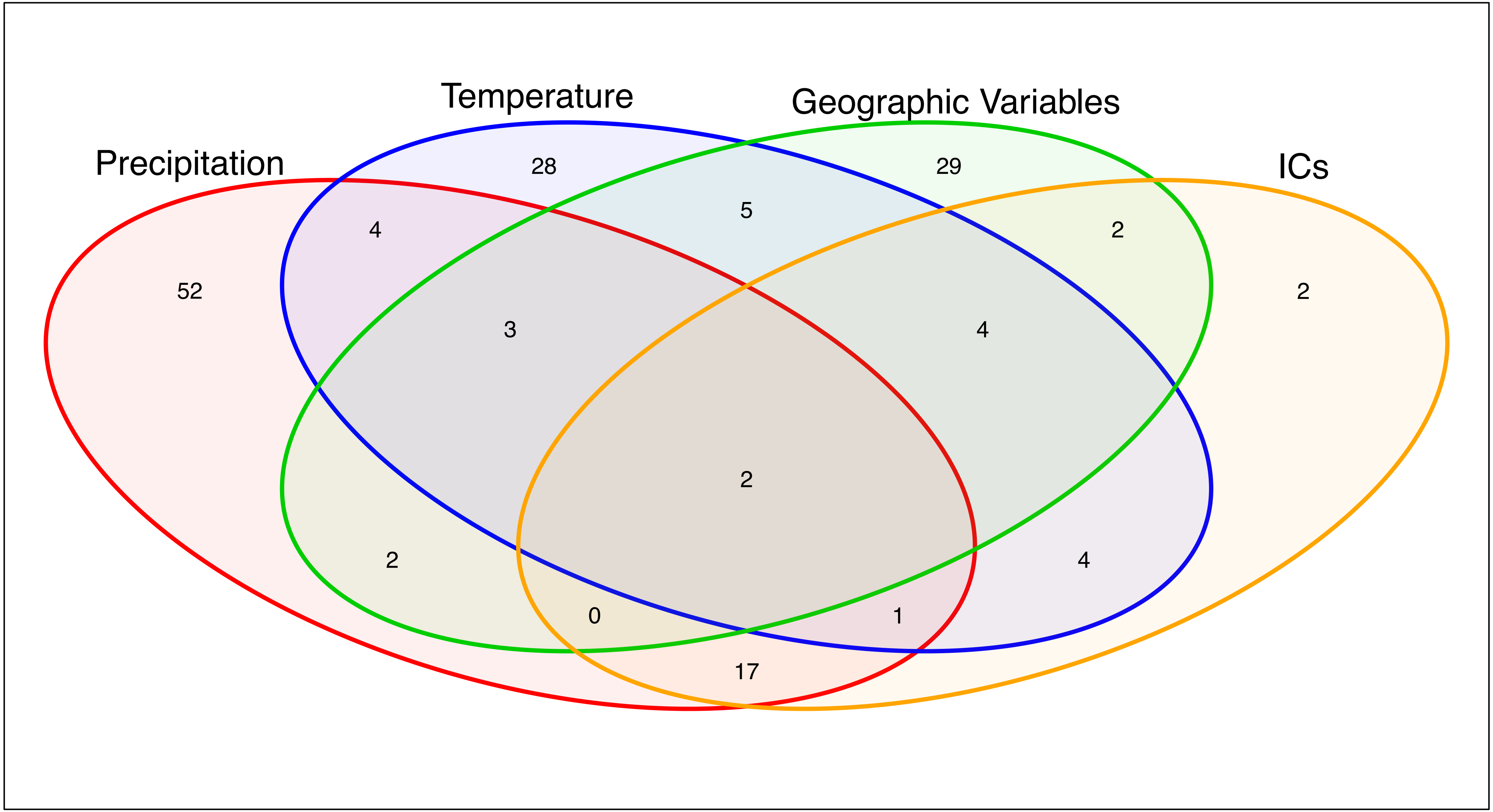
Temperature
Geographic Variables
Precipitation
ICs
28
29
5
4
2
52
2
3
4
2
2
4
0
1
17

**Figure S10**

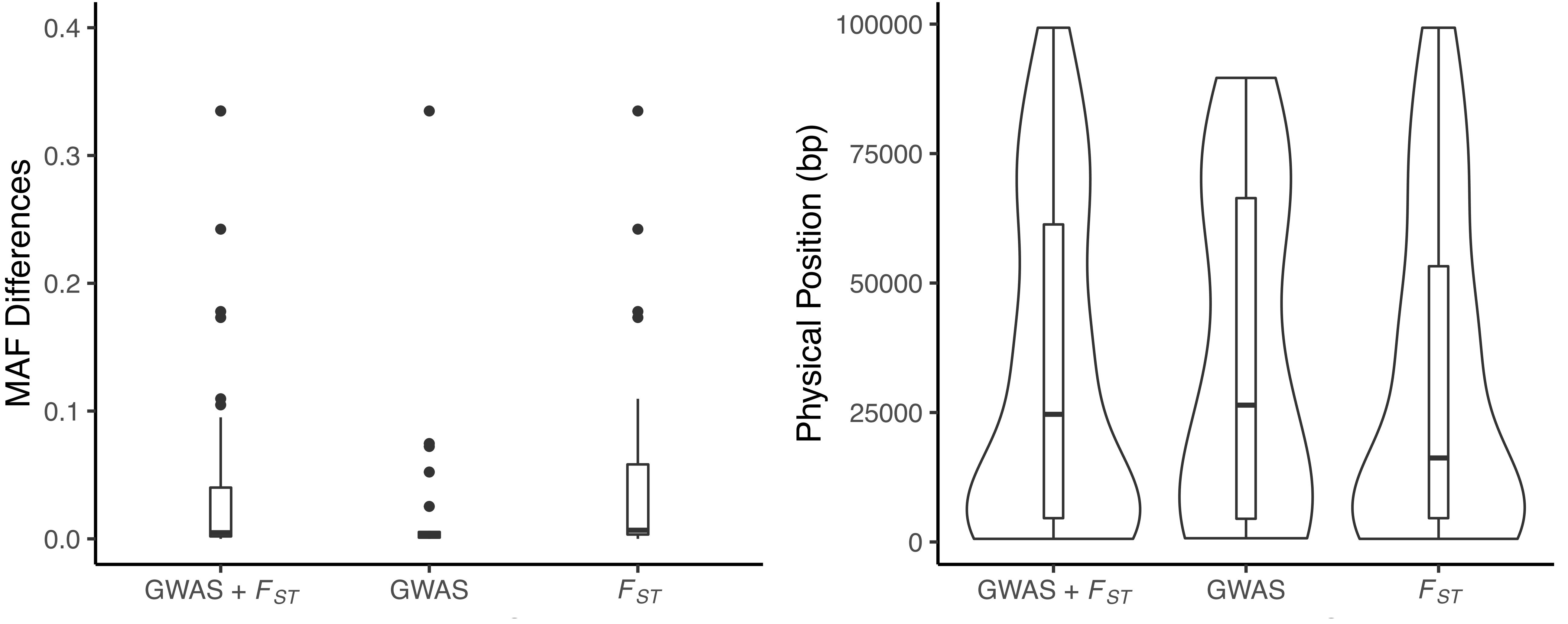
0.4
0.3
0.2
0.1
0.0
MAF Differences
GWAS + $F_{ST}$
GWAS
$F_{ST}$
100000
75000
50000
25000
0
Physical Position (bp)
GWAS + $F_{ST}$
GWAS
$F_{ST}$

**Figure S11**

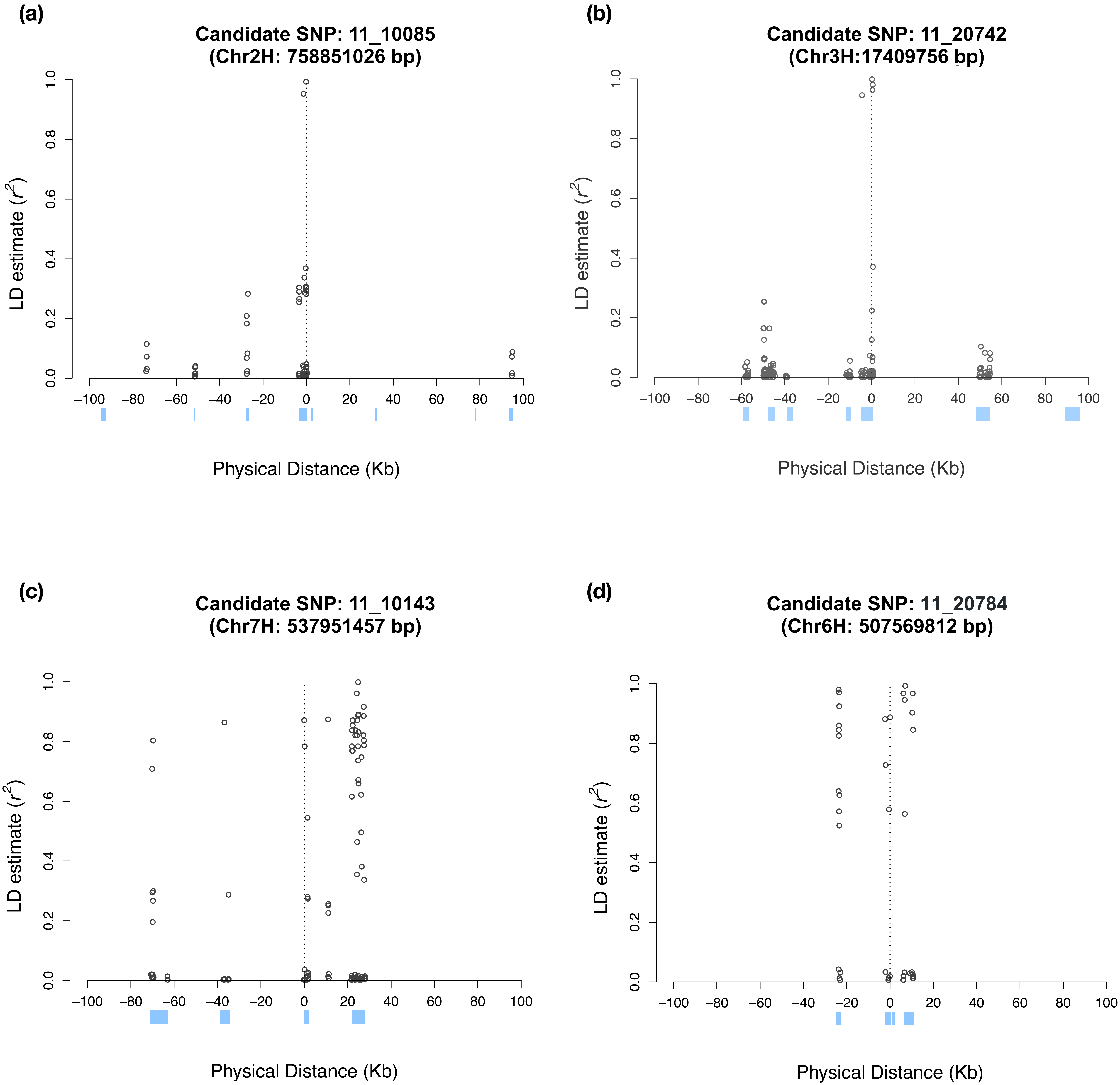
(a)
Candidate SNP: 11_10085
(Chr2H: 758851026 bp)
LD estimate ($r^2$)
Physical Distance (Kb)
(b)
Candidate SNP: 11_20742
(Chr3H:17409756 bp)
LD estimate ($r^2$)
Physical Distance (Kb)
(c)
Candidate SNP: 11_10143
(Chr7H: 537951457 bp)
LD estimate ($r^2$)
Physical Distance (Kb)
(d)
Candidate SNP: 11_20784
(Chr6H: 507569812 bp)
LD estimate ($r^2$)
Physical Distance (Kb)

**Figure S12**

(a)

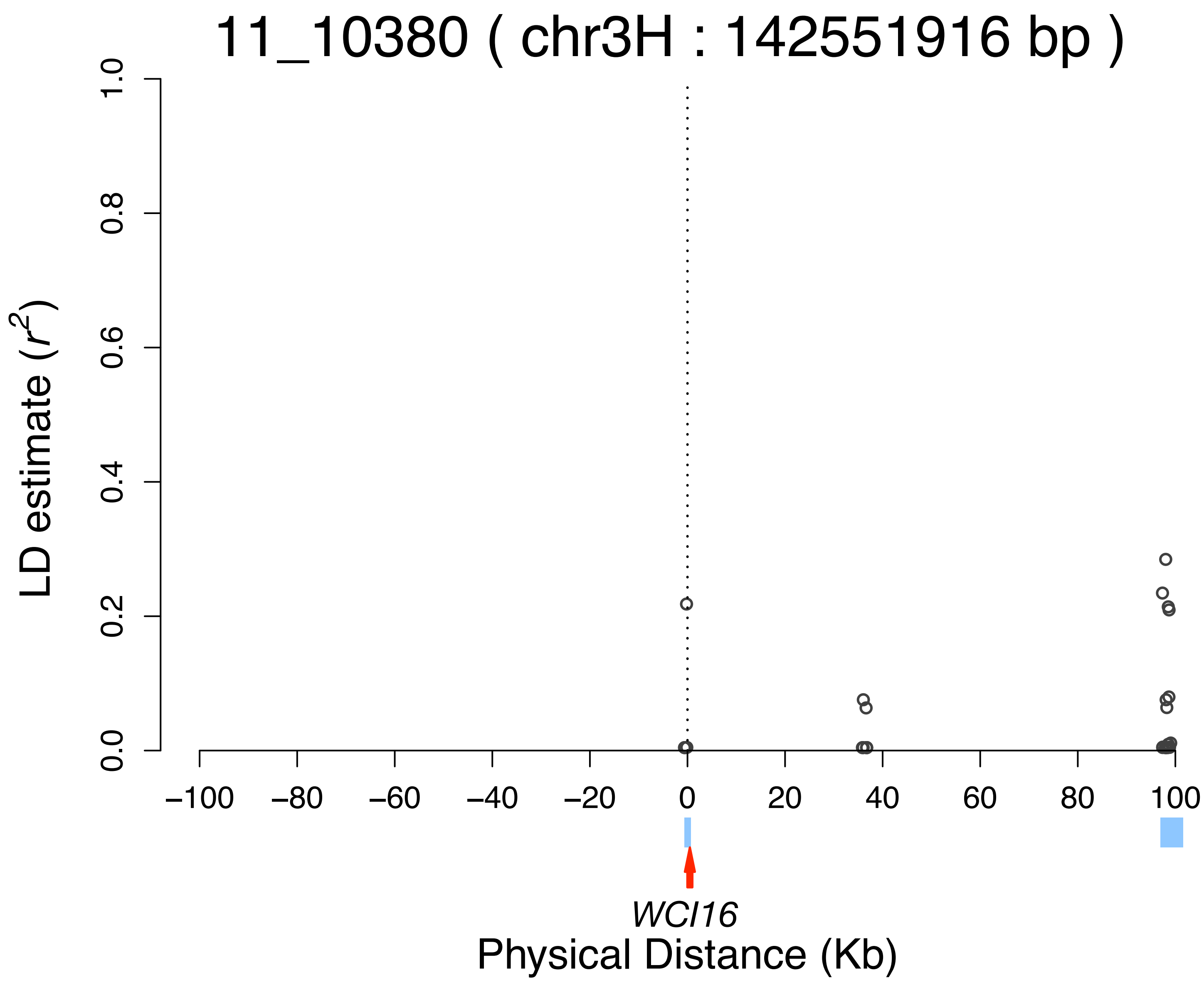


(b)

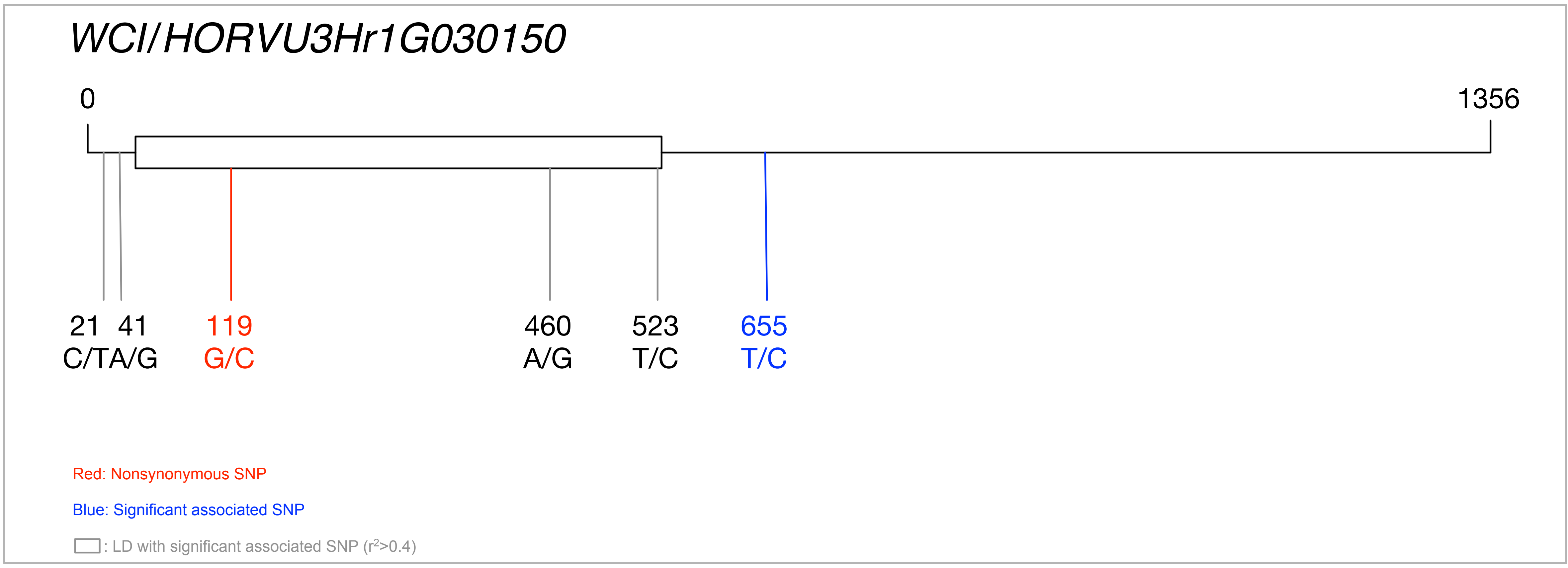


(c)

| Sites | 21 | 41 | 119 | 460 | 523 | 655 | General | | | | Subsets | | | |
|---|---|---|---|---|---|---|---|---|---|---|---|---|---|---|
| Physical position | 1282 | 1302 | 1380 | 1721 | 1784 | 1916 | Elevation | | Latitude | | L_Latitude Elevation | | H_Latitude Elevation | |
| Consensus | C | A | G | G | T | T | | | | | | | | |
| Ancestral | . | . | . | - | . | . | L | H | L | H | L | H | L | H |
| Hap1 | . | . | . | . | . | . | 21 | 2 | 16 | 7 | 14 | 2 | 7 | - |
| Hap2 | . | . | . | . | . | C | 13 | - | 5 | 8 | 5 | - | 8 | - |
| Hap3 | . | . | . | A | . | C | 1 | - | 1 | - | 1 | - | - | - |
| Hap4 | . | . | . | A | . | . | 31 | 2 | 21 | 12 | 19 | 2 | 12 | - |
| Hap5 | . | . | C | . | . | . | 1 | - | 1 | - | 1 | - | - | - |
| Hap6 | . | . | C | A | . | . | 1 | - | 1 | - | 1 | - | - | - |
| Hap7 | . | G | . | A | . | . | 1 | - | 1 | - | 1 | - | - | - |
| Hap8 | T | . | . | . | C | . | 1 | - | 1 | - | 1 | - | - | - |

**Figure S13**

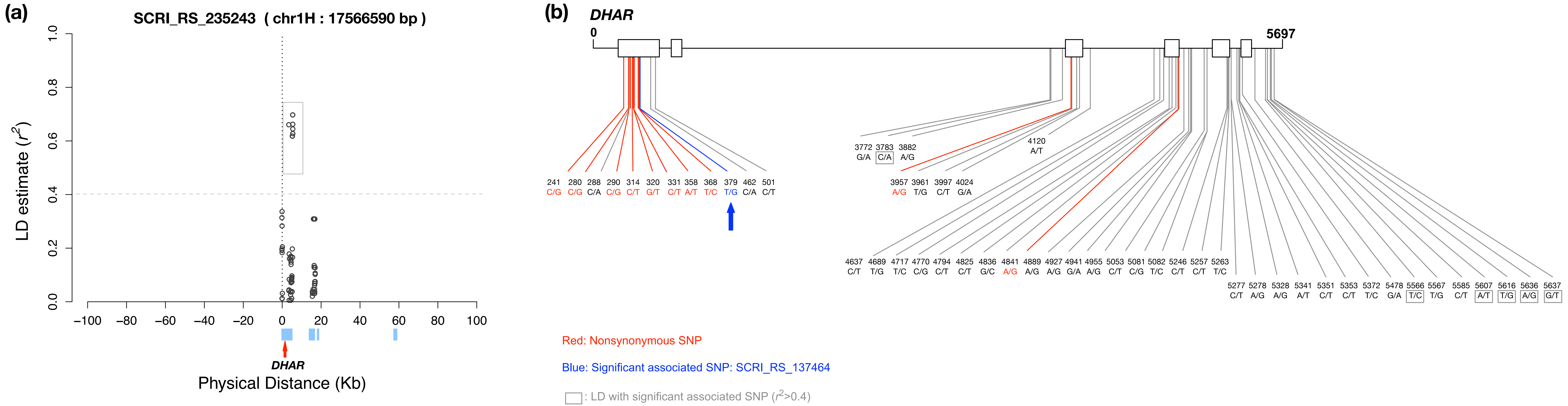


(c)

| Sites | 241 | 280 | 288 | 290 | 314 | 320 | 331 | 358 | 368 | 379 | 462 | 501 | 3772 | 3783 | 3882 | 3957 | 3961 | 3997 | 4024 |
|---|---|---|---|---|---|---|---|---|---|---|---|---|---|---|---|---|---|---|---|
| Physical position | 66452 | 66491 | 66499 | 66501 | 66525 | 66531 | 66542 | 66569 | 66579 | 66590 | 66673 | 66712 | 69983 | 69994 | 70093 | 70168 | 70172 | 70208 | 70235 |
| Consensus | C | C | C | C | C | G | C | A | T | G | C | C | G | A | A | A | T | C | A |
| Ancestral | . | - | . | . | T | . | . | T | . | - | . | . | A | . | . | . | . | . | . |
| Hap1 | . | . | . | . | . | . | . | . | . | T | . | . | . | C | . | . | . | . | G |
| Hap2 | . | . | . | . | . | . | . | . | . | T | . | . | . | C | . | . | . | . | G |
| Hap3 | . | . | . | . | . | . | . | . | . | . | A | . | . | . | . | . | . | T | . |
| Hap4 | . | . | . | . | . | . | . | . | . | . | . | . | . | . | . | . | . | T | . |
| Hap5 | . | . | . | . | . | . | . | . | . | . | . | . | . | C | . | . | . | . | G |
| Hap6 | . | . | . | . | . | . | . | . | . | . | . | T | . | . | . | . | G | . | . |
| Hap7 | . | . | . | . | . | . | . | T | C | . | . | . | . | . | . | . | . | T | . |
| Hap8 | . | . | . | G | . | T | . | . | . | . | . | T | . | . | . | . | . | T | . |
| Hap9 | G | . | . | . | T | T | . | . | . | . | . | . | A | . | G | . | . | . | . |
| Hap10 | G | . | . | . | T | T | . | . | . | . | . | . | A | . | G | . | . | . | . |
| Hap11 | G | G | A | . | T | T | T | . | . | . | . | T | . | . | . | G | . | . | . |

| Sites | 4120 | 4637 | 4689 | 4717 | 4770 | 4794 | 4825 | 4836 | 4841 | 4889 | 4927 | 4941 | 4955 | 5053 | 5081 | 5082 | 5246 | 5257 | 5263 | 5277 | 5278 |
|---|---|---|---|---|---|---|---|---|---|---|---|---|---|---|---|---|---|---|---|---|---|
| Physical position | 70331 | 70848 | 70900 | 70928 | 70981 | 71005 | 71036 | 71047 | 71052 | 71100 | 71138 | 71152 | 71166 | 71264 | 71292 | 71293 | 71457 | 71468 | 71474 | 71488 | 71489 |
| Consensus | A | C | T | T | C | C | C | G | A | A | A | G | A | C | C | T | C | C | T | C | A |
| Ancestral | T | . | . | . | . | . | T | C | . | . | . | A | . | T | . | . | . | . | C | A | . |
| Hap1 | . | . | . | . | . | . | . | . | . | . | . | . | . | . | . | . | . | . | . | . | . |
| Hap2 | . | . | . | . | . | . | . | . | . | . | . | . | . | . | . | . | . | . | . | . | . |
| Hap3 | . | . | . | . | . | . | . | . | . | . | G | . | . | T | G | C | . | . | C | T | G |
| Hap4 | . | . | . | . | . | . | . | . | . | . | G | . | . | T | G | C | . | . | C | T | G |
| Hap5 | . | . | . | . | . | . | . | . | . | . | . | . | . | . | . | . | . | . | . | . | . |
| Hap6 | . | . | . | . | . | . | . | . | . | . | . | . | . | T | G | C | T | T | C | T | G |
| Hap7 | . | . | G | . | . | . | . | . | . | . | G | . | . | T | G | C | . | . | C | T | G |
| Hap8 | . | . | . | . | . | . | . | . | . | . | G | . | . | T | G | C | . | . | C | T | G |
| Hap9 | . | . | . | C | . | T | . | . | . | . | . | . | . | . | . | . | . | . | . | . | . |
| Hap10 | . | . | . | C | . | T | T | . | . | . | . | . | . | . | . | . | . | . | . | . | . |
| Hap11 | T | T | . | . | G | . | . | C | G | G | . | A | G | . | . | . | . | . | . | . | . |

| Sites | 5328 | 5341 | 5351 | 5353 | 5372 | 5478 | 5566 | 5567 | 5585 | 5607 | 5616 | 5636 | 5637 | General: Elevation | | General: Latitude | | Subsets: L_Latitude Elevation | | Subsets: H_Latitude Elevation | |
|---|---|---|---|---|---|---|---|---|---|---|---|---|---|---|---|---|---|---|---|---|---|
| Physical position | 71539 | 71552 | 71562 | 71564 | 71583 | 71689 | 71777 | 71778 | 71796 | 71818 | 71827 | 71847 | 71848 | | | | | | | | |
| Consensus | A | A | C | C | T | G | C | T | C | T | G | G | T | | | | | | | | |
| Ancestral | . | T | . | . | A | . | T | - | . | . | T | A | G | L | H | L | H | L | H | L | H |
| Hap1 | . | . | . | . | . | . | T | . | . | A | T | A | G | 11 | 1 | 8 | 4 | 7 | 1 | 4 | - |
| Hap2 | . | . | . | . | . | . | T | . | C | A | T | A | G | 4 | - | - | 4 | - | - | 4 | - |
| Hap3 | G | T | T | T | C | . | . | . | . | . | . | . | . | 4 | 1 | 4 | 1 | 3 | 1 | 1 | - |
| Hap4 | G | T | T | T | C | . | . | G | . | . | . | . | . | 1 | - | 1 | - | 1 | - | - | - |
| Hap5 | . | . | . | . | . | . | T | . | . | A | T | A | G | 4 | - | 3 | 1 | 3 | - | 1 | - |
| Hap6 | G | T | T | T | C | . | T | . | . | A | T | A | G | 2 | - | 2 | - | 2 | - | - | - |
| Hap7 | G | T | T | T | C | . | . | . | . | . | . | . | . | 4 | - | 3 | 1 | 3 | - | 1 | - |
| Hap8 | G | T | T | T | C | . | . | G | . | . | . | . | . | 2 | - | 1 | 1 | 1 | - | 1 | - |
| Hap9 | . | . | . | . | . | A | . | . | . | . | . | . | . | 5 | 1 | 3 | 3 | 2 | 1 | 3 | - |
| Hap10 | . | . | . | . | . | A | . | . | . | . | . | . | . | 1 | - | 1 | - | 1 | - | - | - |
| Hap11 | . | . | . | . | . | . | . | . | . | . | . | . | . | 21 | - | 13 | 8 | 13 | - | 8 | - |

**Figure S14**
